# CRISPR/Cas9 editing of the wheat iron sensor *TaHRZ1* confirms its conserved role in iron homeostasis and allocation in grains

**DOI:** 10.1101/2025.06.29.662202

**Authors:** Deepshikha Tyagi, Hamida Banoo, Deepak Kumar Jha, Varsha Meena, Riya Joon, Kanupriya Agrwal, Pooja Yadav, Anil Kumar, Santosh B. Satbhai, Terri Long, Ajay K Pandey

## Abstract

Plants rely on specialized sensing systems, including transcriptional regulators, to maintain iron (Fe) homeostasis. Among these, Hemerythrin RING Zinc finger (HRZ) proteins have emerged as key regulators of Fe homeostasis. In this study, six *Triticum aestivum* (wheat) HRZ homoeologs belonging to *TaHRZ1* and *TaHRZ2,* were identified by BLAST search using rice (*Oryza sativa)* HRZ sequences and mapped to chromosomes 1 and 3. These encode proteins with conserved N-terminal Hemerythrin (HHE) domains and C-terminal CHY-RING and Zn-ribbon motifs. Phylogenetic analysis grouped these genes into distinct clades, while expression profiling revealed strong root-specific and Fe-responsive expression patterns, indicating roles in nutrient sensing. Functional conservation was demonstrated by complementation of the *Arabidopsis thaliana bts-1* mutant, where both wheat genes restored normal Fe regulation. Full-length *TaHRZ1* and *TaHRZ2* interacted with members of wheat bHLH IVc transcription factors, while truncated versions lacking the RING domain did not, emphasizing their conserved role in protein interactions. CRISPR-Cas9 editing of the conserved HHE3 domain in all the *TaHRZ1* homoeologs, coupled with GRF4-GIF1 chimeric protein, achieved 6.4-8.8% regeneration efficiency in wheat. Elemental analysis indicated enhanced Fe loading in the grains of the edited lines, particularly in the scutellum, suggesting improved iron partitioning compared to the wild type. Additionally, qRT-PCR revealed upregulation of *TaFIT* and *TaIRO3*, and downregulation of *IDEF1* in edited lines, supporting an important regulatory role for *TaHRZ1* in Fe homeostasis signalling. These findings position *TaHRZ1* as a valuable target for biofortification strategies to enhance Fe content in wheat grains.

## Introduction

Iron (Fe) is among the essential micronutrients in plants that participate as catalytic cofactors in several key processes. Our limited understanding of many of the mechanisms involved in plant Fe homeostasis is a major obstacle to devising approaches for Fe biofortification of staple foods. Plants have developed two strategies to acquire Fe^2+^ from the rhizosphere (Kobayashi & Nishizawa 2012). Non-graminaceous plants utilize a reduction-based strategy (Strategy I), while gramineous plants, which include major cereal staple crops, utilize a chelation-based strategy (Strategy II). The Fe uptake process in Strategy I involves the release of protons into the rhizosphere by H^+^-ATPases, resulting in acidification and increased Fe solubility (Santi & Schmidt 2009). Subsequently, ferric (Fe^3+^) is reduced to ferrous (Fe²D) by ferric chelate reductases (FCR). In *Arabidopsis thaliana,* FRO2 (a membrane-bound FCR) is one of the eight members of the FRO protein family that serves this function (Robinson, Procter, Connolly & Guerinot 1999). In Arabidopsis, lastly, Fe^2+^ is translocated across the plasma membrane into the cells by an Fe-regulated transporter (IRT1), a membrane-bound transporter that is a member of the zinc-regulated transporter (ZRT), IRT-like protein (ZIP) divalent metal transporter family (Connolly, Fett & Guerinot 2002). Strategy I plants, such as *Arabidopsis thaliana*, also release compounds like catechol coumarins that facilitate Fe uptake by enhancing Fe³D reduction in the rhizosphere (Schmid et al. 2014; Rajniak et al. 2018). In contrast, bread wheat is a clear Strategy II plant, acquiring Fe exclusively via the chelation-based pathway. Strategy II involves the biosynthesis and release of mugineic acid family phytosiderophores (PS) into the rhizosphere. Fe is then solubilized through complex formation with PS, and Fe^3+^-PS complexes are transported into the root via transporters belonging to the Yellow-Stripe (YS) family of proteins (Curie et al. 2001; Palmer & Guerinot 2009) .

In addition to Fe, IRT1 also transports other metal ions such as Zinc (Zn), Manganese (Mn), and Cadmium (Cd). (Eide, Broderius, Fett & Guerinot 1996; Vert et al. 2002; Colangelo & Guerinot 2004). Once absorbed, Fe binds to chelating molecules like nicotianamine and citrate, which assist in its transport into and throughout the plant’s vascular system (Durrett et al., 2007; Curie et al., 2009). At the transcriptional level, a major transcription factor FER-like Fe Deficiency-Induced Transcription Factor (FIT) in *Arabidopsis thaliana* regulates the expression of various Fe uptake genes (Bauer, Ling & Guerinot 2007). Additionally, other regulators, especially the basic helix-loop-helix (bHLH) transcription factors, including bHLH38, bHLH39, bHLH100, and bHLH101, interact with FIT to enhance its activity (Yuan et al. 2008). Thus, FIT acts as a regulatory hub and forms complexes with other proteins to activate target genes involved in Fe homeostasis.

Ubiquitination mediated by E3 Ubiquitin ligases, has been shown to be an important mechanism for controlling the turnover of Fe transporters and transcription factors (TFs) (Barberon et al., 2011; Dubeaux et al., 2018; Maria N. Hindt et al., 2017; Kobayashi et al., 2013; Spielmann and Vert, 2021). Specifically, Hemerythrin-rich Really Interesting New Gene (RING)- and Zinc-finger (HRZ) proteins in rice (*Oryza sativa* )and its *Arabidopsis* homolog BRUTUS (BTS) and BRUTUS-like (BTSL1 and BTSL2) play a significant role in maintaining Fe homeostasis in plants (Rodríguez-Celma et al., 2019; Gao et al., 2020). HRZ proteins form a subfamily of hemerythrin E3 ubiquitin ligases that are conserved from green algae to higher plants and bear distant homology to mammalian F-box and Leucine-rich Repeat Protein 5 (FBLX5), a post-translational regulator of Fe homeostasis (Salahudeen et al. 2009; Vashisht et al. 2009). HRZ/BTS proteins are characterized by their combination of hemerythrin (HHE) domain repeats, which can bind Fe, and RING and zinc-finger motifs, commonly associated with protein-protein interactions and C-terminal ubiquitin ligase activity (Selote et al., 2015; Rodriguez-Celma et al.,2019).

In rice two HRZ homologs (HRZ1 and HRZ2) were reported and specifically, *OsHRZ1* was shown to affect the stability of bHLH transcription factor members involved in Fe uptake through its E3 ubiquitin ligase activity (Kobayashi et al. 2019). In general, HRZ1/BTS proteins possess three HHE domains at the protein level, while BTSL proteins have only two HHE domains (Maria N. Hindt et al., 2017; Kobayashi et al., 2013) . Previous work has demonstrated that the deletion of the HHE domain does not impact the polyubiquitination activity of HRZ proteins, indicating that the ubiquitination function is conserved even in the absence of this domain (Kobayashi et al., 2013) However, removing the C-terminal domains, which included the RING domain, eliminated the ubiquitination activity. It has been speculated that, while the HHE domain does not affect ubiquitination activity, it could contribute to the overall function of HRZ proteins by enabling them to respond to metal status or affecting their regulatory roles in Fe homeostasis (Kobayashi et al., 2013). At the functional level, BTS lacking the HHE domain could complement the *bts-1* sensitive phenotype, suggesting its contribution to the instability of the protein (Selote, et al., 2015). Similarly, IRT1 was also shown to undergo ubiquitin-conjugate-mediated degradation mediated by Iron Degradation factor RING E3 ligase (IDF1) (Dubeaux et al., 2018; Shin et al., 2013). In *Arabidopsis thaliana*, BTS mediates the ubiquitination and degradation of Fe-related bHLH transcription factors such as FIT and subgroup Ib bHLHs, whereas in *O. sativa*, HRZ proteins regulate iron homeostasis by targeting IRO/PRI transcription factors, highlighting species-specific downstream targets of the HRZ/BTS pathway. These studies underscore the critical role of E3 ubiquitin ligases in regulating iron homeostasis and highlight the need for further investigation in important crop species.

Hexaploid wheat is a major global crop and a key target for micronutrient biofortification. Previously, our group identified a comprehensive set of Fe-responsive genes in wheat and uncovered several potential regulatory candidates (Kaur et al. 2019). Functional assessment of wheat genes shall enable bioengineering key regulators to drive targeted strategies for grain Fe biofortification. For this, efficient genetic transformation in wheat remains a major bottleneck due to strong genotype dependency and the limited regeneration capacity of many elite cultivars in tissue culture. To overcome the recalcitrance of wheat to tissue culture and genetic transformation, several developmental regulators have recently been explored to enhance regeneration efficiency. Growth Regulating Factor 4 (GRF4), aided by its cofactor GRF-Interacting Factor (GIF1), enhances wheat transformation efficiency by promoting cell proliferation and regeneration competency in tissue culture (Debernardi et al. 2020).

In the current study, we identified and characterized wheat homologs of rice HRZ, referred to as *TaHRZ*1 and *TaHRZ*2, which encode conserved domain structures and exhibit strong root-specific and Fe deficiency-induced expression. Functional complementation in Arabidopsis and interaction with bHLH IVc TFs support a conserved role in Fe signalling. Importantly, CRISPR-Cas9-mediated editing of *TaHRZ*1, aided by GRF4 and its GIF1, enhanced grain Fe content without yield penalties under control conditions. These findings highlight *TaHRZ*1 as a promising target for wheat biofortification.

## Material and methods

### Plant materials and growth conditions

Hexaploid wheat cv. C-306 and Fielder were used for this study. Following an overnight stratification process at 4 °C in the dark, wheat seeds were allowed to germinate on Petri dishes lined with Whatman filter paper. Endosperms were removed from 4 days old seedlings and transferred in PhytaBoxTM containers using Hoagland’s nutritional solution. Plants were grown under −Fe (1 µM Fe-EDTA) or +Fe (80 µM Fe-EDTA) conditions, without any changes to the amount of other nutrients. The plants were grown in a controlled growth environment with a photon flux density of 300 µmol m^−2^ s^−1^, at 21±1°C, 50–65% relative humidity, and a 16–8hour light-dark photoperiod. After eight days under −Fe and +Fe conditions, roots and shoots were collected from three biological replicates and promptly frozen in liquid nitrogen for further examination. Iron concentrations (1 µM and 80 µM Fe-EDTA) were selected based on previous wheat Fe-deficiency studies and preliminary optimization experiments (Kaur et al. 2019; Meena et al. 2024).

*Arabidopsis thaliana* ecotype Columbia (Col-0) was used as wild type (WT) along with *atbts1* mutant. Arabidopsis seeds were surface sterilized by 2% sodium hypochlorite for 5 minutes, followed by wash with 70% ethanol for 2 minutes, and subsequently washed with autoclaved water for 5 times. The sterilized seeds were then stratified for 3 days. Seeds were then germinated and grown on Hoagland medium for 5 days under control conditions to establish uniform growth. Then seedlings were moved to minimum media that had been altered with + Fe (50 µM Fe-EDTA; control) and − Fe (1 µM Fe-EDTA supplied with 200 µM ferrozine; deficiency) for an additional 7 days.

After treatment, roots and shoots were collected from three biological replicates and promptly frozen in liquid nitrogen for further examination.

### Phylogenetic study and in-silico identification of wheat HRZ genes

HRZ protein sequences from *O. sativa*, BTS from *A. thaliana* (the hemerythrin-domain E3 ligase), and its two homologs (TaHRZ1 and TaHRZ2) from *Triticum aestivum* were utilized for phylogenetic analysis. *A. thaliana* protein sequence was used to retrieve these HRZ protein sequences from the entire wheat genome using BLAST search in the EnsemblPlant. Additionally, BLAST using rice (*Oryza sativa*) HRZ gave higher similarity, reflecting the closer rice–wheat relationship. The MUSCLE algorithm (Edgar 2004) was used to align the putative IDs and protein sequences, and MEGAX software was used to create a rooted tree using the neighbour joining method (Kumar, Stecher, Li, Knyaz & Tamura 2018). The phylogenetic tree was created to show the HRZs of *O. sativa* and *Arabidopsis BTS*, which are most related to wheat HRZ. The conserved motif architecture of the TaHRZ transcripts was subsequently ascertained by analysing their gene structure. MEME was used to anticipate these conserved motifs(Bailey, Johnson, Grant & Noble 2015). TaHRZ and Arabidopsis both shared conserved N-terminal Hemerythrin (HHE) domains and C-terminal CHY-RING and Zn-ribbon motifs. Additionally, the Gene Structure Display Server (GSDS) was used to analyse the organization of the introns and exons of wheat HRZ and their homeologs (Hu et al. 2015).

### Arabidopsis thaliana complementation and experimental conditions

Using site-directed cloning, the *TaHRZ*1 (ORF: 3714 bp; *TraesCS3A02G262700*) and *TaHRZ*2 (ORF: 3645 bp; *TraesCS1A02G374800*), which were strongly expressed, were cloned into the plant expression vector pCAMBIA1302 (pCAMBIA1302:*TaHRZ*1/*TaHRZ2*-His) at the *Bgl*II and *Spe*1 restriction sites. These constructs were introduced into wild type and *atbts1*(SALK_016526) mutant plants of *A. thaliana*, using *Agrobacterium tumefaciens* (GV3101) mediated transformation by the floral dip method (Zhang, Henriques, Lin, Niu & Chua 2006). Multiple (7-10) independent transformants were screened on 0.5x MS media containing 30 mg/L hygromycin and 0.8 % agar. The transformed seedlings (T_1_) with long hypocotyls and green expanded leaves at a 4-leaf stage were separated from the non-transformed seedlings and transferred to the soil after about 3 weeks. Similarly, T_2_ and T_3_ generation seeds were also selected and allowed to grow till maturity. The transgenic seedlings were confirmed for the presence of recombinant cassette using PCR-based approach. At least three independent homozygous T_3_ lines per construct were used for phenotypic, molecular, and biochemical analyses. The transgenic lines containing the empty pCAMBIA1302 vector were used as a vector control. The PCR-positive lines, which showed Western blot positivity, were further used for functional characterisation.

### Design of guide RNA, wheat transformation, screening of TaHRZ1 edits and genotyping

The conserved region of all three homoeologs of *TaHRZ*1 was used for gRNA design. The gRNA spacers were designed using wheat CRISPR webtool (Breaking Cas software https://bioinfogp.cnb.csic.es/tools/breakingcas/). We designed the gRNA to target a unique site in the 7^th^/8^th^ exon of the *TaHRZ*1 gene that was a consensus site for all three homoeologs (**Table S1**). The *TaHRZ1* targeting gRNA construct prepared in JD633 vector was introduced into *Agrobacterium* strain AGL-1 and screened. cv. ‘Fielder’ and C-306 were transformed using *Agrobacterium*-mediated transformation, as described earlier (Bhati et al., 2016; Hayta et al., 2021). Hygromycin was used for the selection of transformed calli at different stages of tissue culture: 15 mg/L during the initial callus selection, 30 mg/L during subsequent callus proliferation, and 20 mg/L during shoot regeneration.

The presence of the transgene in regenerated plants was confirmed by PCR analysis using multiple pairs of Hygromycin cassette-specific primers, following the methodology described by (Bhati et al., 2016). Genomic DNA was extracted using the DNeasy Plant Pro Kit (Qiagen, Hilden, Germany) from leaf discs collected from plants carrying the same constructs, following the manufacturer’s protocol with the exception of using double the volume of grinding buffer. Target regions were amplified using conserved primers (**Table S1**) and GoTaq DNA Polymerase (Promega, USA) according to the manufacturer’s instructions. A ∼350 bp amplicon was generated from genomic DNA extracted from the edited lines. Firstly, these amplicons were subjected to the T7 Endonuclease *I* (T7E1) assay as described earlier (Kim, Ramakrishna, Kim & Kim 2014). To verify the occurrence of a DNA double-strand break, PCR amplicons were purified from agarose gel, denatured, reannealed, and subsequently T7EI assay was performed. The amplicons showing fragmentation were subjected to next-generation sequencing (Eurofins, Bangalore, India). Mutagenesis in specific genomic regions was confirmed using Integrative Genomics Viewer (IGV) (Thorvaldsdóttir, Robinson & Mesirov 2013). Additionally, Sanger sequencing (Eurofins Scientific DNA Sequencing Core, Bangalore, India) of the independently cloned PCR fragments was conducted to validate the nature of the edits as reflected in NGS data. Resulting sequences were analysed using the Synthego ICE Analysis tool, which compares mutant sequences against the wild-type reference to determine editing efficiency or knockout scores. Only edited lines showing consistent indel profiles across technical replicates were selected for downstream phenotypic and elemental analyses.

### Perls staining and elemental analysis

Root samples from the above experiments were used for Perl’s Prussian Blue staining to evaluate Fe^3+^ accumulation. Root samples were vacuum infiltrated for an hour in a dark environment to improve reagent penetration after being submerged in a solution containing 5% potassium ferrocyanide and 5% HCl. Following staining, the roots were carefully cleaned with deionized water to remove any remaining stains examined under a stereomicroscope. The development of a blue precipitate indicated the presence of Fe³^+^. For elemental analysis, pooled harvest of tissues was done to collect samples from *A. thaliana* or wheat seedlings to minimize intra-plant variation. The samples were oven-dried, digested in nitric acid and hydrogen peroxide, and analyzed for Fe content using an ICP-MS instrument (Agilent 7900, USA) following standard protocols described earlier (Meena et al. 2024; Agrwal, Veena, Singhal & Pandey 2025). Data are expressed as mg Fe kg^−1^ dry weight. Similarly, Zn (Zinc), Copper (Cu) and Manganese (Mn) concentrations were measured in the digested samples.

### Isolation of total RNA, cDNA synthesis, and quantitative real-time PCR analysis

RNA was manually isolated from diverse tissues using TRIzol® Reagent (Invitrogen™). To remove genomic DNA contamination, RNA samples were treated with TURBO™ DNase (Ambion, Life Technologies) after quantifying their integrity and concentration. Two micrograms of total RNA were used to synthesize cDNA as per the manufacturer’s instructions using the Invitrogen SuperScript III First-Strand Synthesis System SuperMix (Thermo Fisher Scientific). QuantiTect SYBR Green RT-PCR Kit (Qiagen, Germany) was used for qRT-PCR. Homoeolog-non-discriminating primer sets targeting conserved regions of *TaHRZ1* and *TaHRZ2* were used to amplify transcripts from all three sub-genomes representing the additive expression responses. The experiment was validated using four technical replicates for each pair of primers and at least two to three experimental replicates. As reported earlier, for normalization, the ADP-ribosylation factor gene (*TaARF*) was used as an internal control for wheat samples (Kaur et al. 2019), and *AtActin* was used as an internal control for Arabidopsis samples. The stability of these reference genes across tissues and Fe treatments was verified prior to analysis. The C_t_ values obtained after the run were normalised against the internal control, and relative expression was assessed using the 2^−ΔΔCT^ method (Livak & Schmittgen 2001). All the primers used in the study are mentioned in **Table S1**.

### Physiological and molecular analyses of edited lines

Physiological and morphological traits, including plant height, number of grains per line, and 1000-grain weight, were recorded at the maturity stage (T_1_ seeds). Each trait was evaluated using a minimum of three biological replicates from both transgenic edited lines and control plants. Mature grain samples from wheat edited lines and wild type were subjected to Perl’s Prussian Blue staining to visualize Fe³D accumulation, and iron content was quantified using ICP-MS after acid digestion of pooled mature grains. Additionally, wheat seedlings from edited and control lines grown under +Fe and –Fe conditions were used for iron responsive gene expression analysis through qRT-PCR.Seeds post twenty one days anthesis were collected from wild-type and edited lines for expression analysis of transcript encoding the two Ferritin homoeologs *TaFER1*:(*TraesCS5A02G152400*, *TraesCS5B02G151000*, *TraesCS5D02G157600*) and *TaFER2*:(*TraesCS4D02G131700*, *TraesCS4B02G137200*). Five seeds from each edited lines and wild-type were pooled for RNA isolation. cDNA synthesis and qRT-PCR was done as mentioned above.

### Yeast-two-hybrid assays and Bimolecular fluorescence complementation (BiFC) assay

Following PCR amplification, the complete coding sequences of *TaHRZ*1 and *TaHRZ*2 was cloned into the GAL4 activation domain vector pGADT7 (prey) and the GAL4 DNA-binding domain vector pGBKT7 (bait), respectively. To examine domain-specific protein interactions, truncated TaHRZ1 and TaHRZ2 (t-TaHRZ1/2) lacking the C-terminal RING and zinc-finger domains but retaining the N-terminal hemerythrin domains were generated to test domain-specific interactions. These truncated variants were employed to assess the requirement of the RING and zinc-finger domains for interactions with TabHLH IVc and TaFIT transcription factors. The lithium acetate/PEG technique was used to co-transform the yeast strain AH109 (Clontech Matchmaker Gold System, USA) with bait and prey constructions in accordance with the manufacturer’s instructions. The presence of both plasmids was verified by selecting transformants on synthetic dropout (SD) medium deficient in leucine and tryptophan (SD/-Leu/-Trp). Overnight growth of single colonies from SD/-Leu/-Trp plates was conducted at 30°C and 200 rpm in SD/-Leu/-Trp liquid medium. Serial 10-fold dilutions from 10□ to 10□³ were made in sterile distilled water after cultures were adjusted to an OD□□□ of 1.0. The yeast cultures were spotted (5 µL each) onto SD/-Leu/-Trp/-His + X-α-GAL plates in serial 10-fold dilutions from 10□ to 10□³ and incubated for 3–5 days at 30°C. The development of the blue colour and yeast growth were used to evaluate protein–protein interactions.

For BiFC assay, the full-length coding sequences of *TaHRZ1*, *TaFIT*, *TabHLH406*, and *TabHLH408* were cloned into the pSYNE and pSYCE vectors to generate N-terminal (nYFP) or C-terminal (cYFP) fusion proteins, respectively. The resulting constructs were transformed into *A. tumefaciens* GV3101. *Agrobacterium* cultures containing each construct were then used to infiltrate *Nicotiana benthamiana* leaves. The infected leaves were imaged after 48=h using a Leica SP8 confocal laser scanning microscope (Leica Microsystems, Buffalo Grove, IL, USA).

### Phytic acid estimation

Phytic acid (PA) content was quantified using the Megazyme Phytic Acid Assay Kit (K-PHYT, Ireland) following the manufacturer’s protocol and as previously earlier (Agrwal et al. 2025). Briefly, 100 mg of finely ground mature grains was extracted with 2 ml of 0.66 M HCl and incubated overnight at 28 °C with shaking (200 rpm). After centrifugation at 13,000 rpm, 0.5 ml of the supernatant was collected and neutralized with an equal volume of 0.75 M NaOH. Following neutralization, the determination of free phosphorus and total phosphorus was carried out according to the enzymatic assay procedure provided in the kit. Measurements were done using three independent extractions with three technical replicates for digestions. Phytic acid content was then estimated based on the difference between these values using the Mega-Calc™ software tool supplied by the manufacturer.

### Statistical analysis

Statistical analyses were performed using one-way or two-way ANOVA followed by Tukey’s HSD post hoc test using SPSS software. Data are presented as mean ± standard error (SE) based on at least three independent biological replicates. Normality and homogeneity of variance were verified prior to analysis.

## Results

### Identification, sequence and expression analysis of wheat HRZ genes

To identify candidate TaHRZ proteins in hexaploid wheat (*T. aestivum*), we used *A. thaliana* BTS and rice (*O. sativa) HRZ* protein sequences to perform a BLAST search against the EnsemblPlants wheat genome database. Our analysis identified two paralogous genes on chromosomes 1 and 3, designated as *TaHRZ*1 and *TaHRZ*2, each with three homoeologs from the A, B, and D subgenomes of hexaploid wheat. The full-length cDNAs of *TaHRZ1 and TaHRZ2* were 3732 and 3645 bp, respectively and were expected to encode proteins of 1237 and 1214 aa with >98% identity among homoeologs. These proteins contain three N-terminal HHE domains, and the C-terminal with the presence of zf-CHY, RING-UB, and Zn-ribbon domains (**Figure 1A, Figure S1**).). TaHRZ1 shared 58% identity with BTS, and 78%, 77% with OSHRZ1 and OsHRZ2. Whereas the TaHRZ2 shared 53% identity with BTS, and 73% ,79% with OSHRZ1 and OsHRZ2. BLAST searches using the rice HRZ sequences yielded higher similarity scores, consistent with the closer evolutionary relationship between rice and wheat, whereas inclusion of the *A. thaliana* BTS sequence allowed identification of conserved features across dicot and monocot lineages. The degree of sequence similarity between TaHRZ proteins led to their classification into four clades. TaHRZ proteins resemble *O. sativa* the most, followed by *Arabidopsis*. A Neighbour-joining phylogenetic tree constructed using TaHRZ1 and TaHRZ2 and their homologs from *Arabidopsis thaliana*, *Oryza sativa* grouped the wheat genes into two distinct clades (1^st^ and 2^nd^ clades) (**Figure 1B**). HRZ homologs from diploid progenitors (*Triticum urartu*, A genome; *Aegilops tauschii*, D genome), tetraploid wheat (AABB), and hexaploid bread wheat (AABBDD) clustered according to their genomic origins, confirming the presence of homoeologs in all three sub-genomes (**Figure S1B**). Furthermore, the multiple sequence alignment demonstrates strong conservation of the characteristic N-terminal HHE domains and the C-terminal ZF-CHY/RING-type zinc finger motifs between wheat TaHRZ proteins and their orthologues in *Arabidopsis* and rice. The preservation of these key iron-sensing and ubiquitin ligase–associated domains further supports the evolutionary conservation of HRZ/BTS function across monocot and dicot plant species (**Figure S2**). Expression analysis of *TaHRZ1* and *TaHRZ2* at the homoeolog-specific level revealed differential but tissue-specific patterns. In spike, stem, and root tissues, all three homoeologs (A, B, and D) of both genes were expressed, with markedly higher transcript levels detected in roots compared to spike and stem (**Figure 1C**). Enriched expression in the root suggests a conserved and potentially critical role of *TaHRZs*.

**Figure 1:**
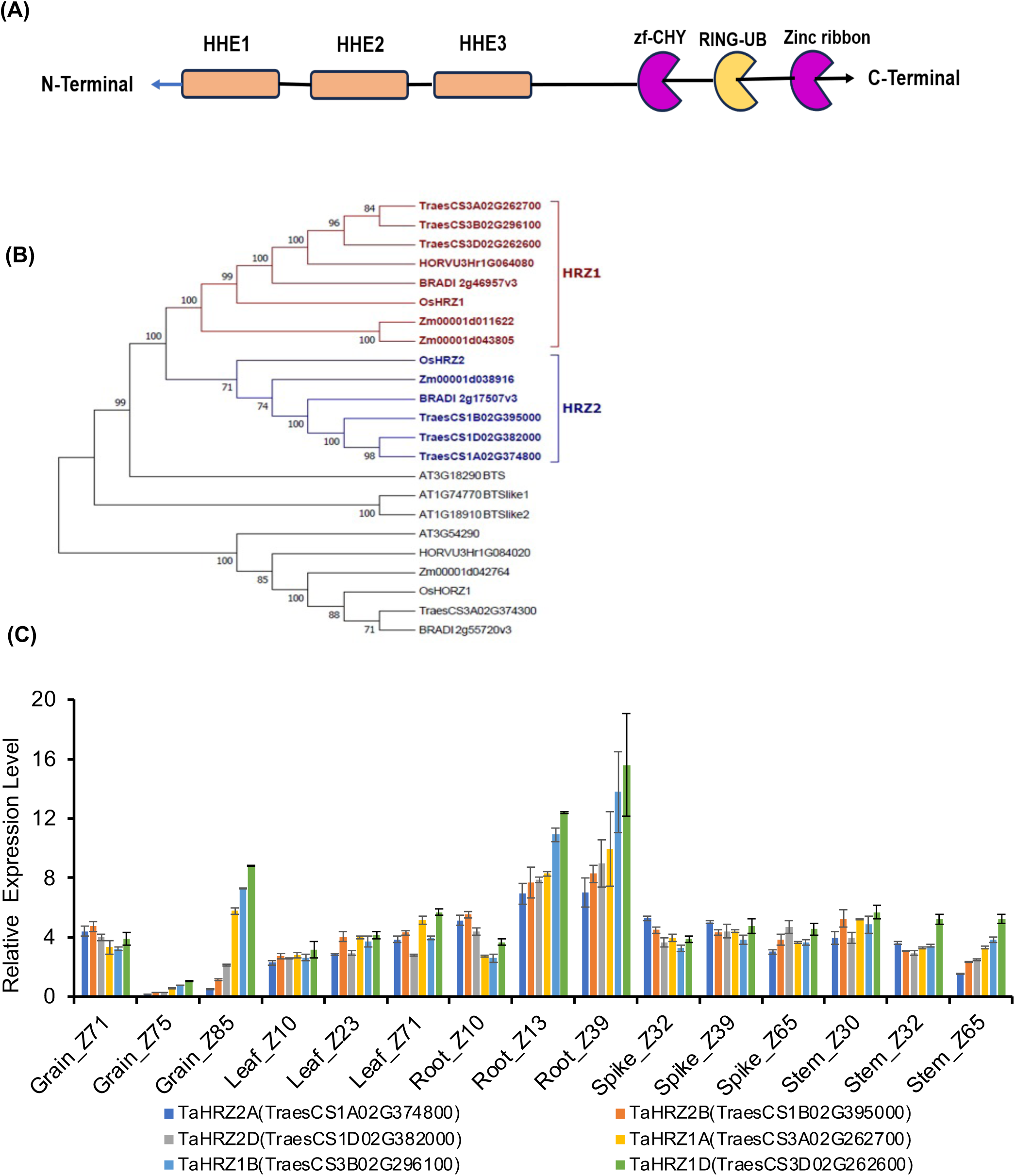
Phylogeny and characterization of *Triticum aestivum* (wheat) HRZ1 and HRZ2 proteins. (A) Schematic representation of the wheat HRZ protein domains. Both proteins contain three Hemerythrin (HHE) domains and a single zf-CHY, RING-UB domain with Zinc ribbon motifs. (B) Phylogenetic analysis of TaHRZ1 and TaHRZ2 homoeologs and other known HRZ proteins from *Arabidopsis* (At), *Zea mays* (Zm), *Brachypodium* (Bd) and rice (Os) homologs grouped them into 1st and 2nd clade main clades. The MUSCLE algorithm was used to align MCO proteins, and the MEGA X neighbour-joining method was used to create an unrooted phylogenetic tree with a bootstrap value of 1000. (C) In-silico expVIP expression analysis of *TaHRZ* gene homoeologs in different wheat tissues at different phases of wheat development.

To gain insight into *TaHRZ1* and *TaHRZ2* transcript accumulation, we examined their gene expression in root and shoot tissues at different time points in response to Fe deficiency (1 µM Fe-EDTA). After 4 days of Fe deficiency, in roots, both *TaHRZ1* and *TaHRZ2* expression were markedly higher than that under +Fe, whereas Fe resupply (R) decreased expression of *TaHRZ2*. (**Figure 2A,B**). In shoots, at 4 D higher fold expression of *TaHRZ1* and *TaHRZ2* was observed as compared to roots (**Figure 2B**). Expression was normalized to TaARF, which was stable across tissues and stages, ensuring comparability. In addition to this, the expression of *TaHRZ*s in wheat grains and tissue type suggested that *TaHRZ*1 is highly expressed at 21 days after anthesis indicating tissue- and stage-specific regulation. (**Figure 2C**). Additionally, both the *TaHRZ* genes were highly expressed in the pericarp region of the grain tissue (**Figure 2D**).

**Figure 2:**
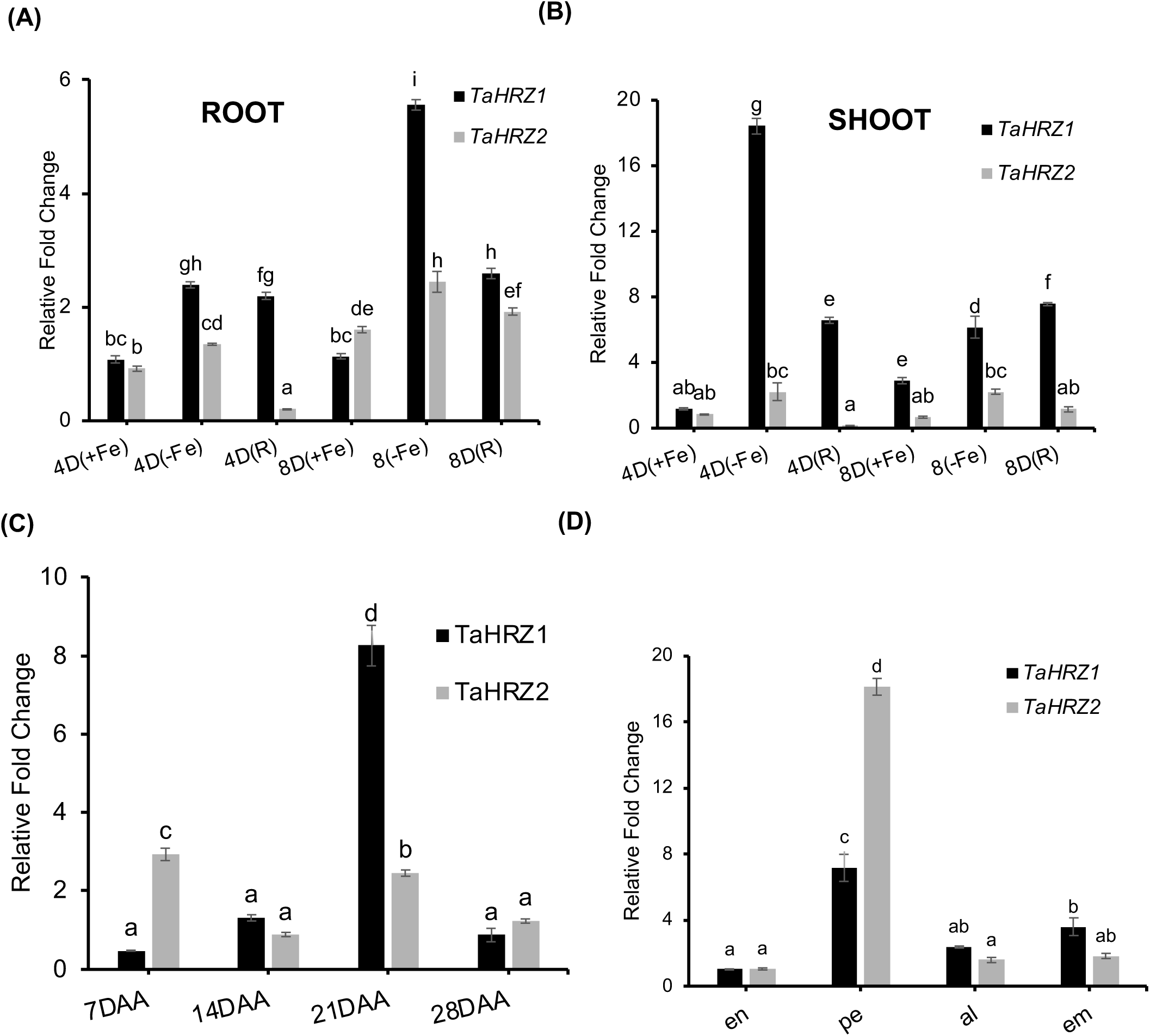
Relative expression analysis of *TaHRZ*1 and *TaHRZ*2 in wheat roots and shoots under Fe deficiency conditions. (A) Relative expression of *TaHRZ*1 and (B) *TaHRZ*2 at 4D and 8D Fe resupply treatment. Fe was supplied as 1 µM Fe-EDTA (−Fe) or 80 µM Fe-EDTA (+Fe). (C) *TaHRZ*1*/HRZ*2 expression in wheat plants varies by different developmental stages. The analysis was occurred at four different developmental stages: 7, 14, 21, and 28 days after anthesis (DAA). (D) Quantitative Expression Analysis of *TaHRZ1/HR*Z2 in several tissues inside the grain, including the p:pericarp; al:aleurone; em:embryo; en:endosperm. To normalize the expression level, TaARF1 was used as an internal reference gene. Fold change were determined using the 2^−ΔΔCT method in all the panels. The standard deviation was calculated from three biological replicates (n=3). Significant difference (P < 0.05) were shown by different letters above the bars using one way Anova analysis.

### Wheat HRZs complement bts-1 and interact with bHLH IVc member proteins

Next, we checked the ability of both the wheat HRZs to rescue the *A. thaliana* BTS function. Arabidopsis overexpressing *TaHRZ1* and *TaHRZ2* in the Arabidopsis *bts-1* mutant were generated, and multiple lines were confirmed for 6XHis-tagged fused HRZ protein by Western (**Figure S3**). At the phenotypic level, under +Fe, the multiple complemented lines (Ox-HRZ1, and Ox-HRZ2), Col-0 and *bts-1,* exhibited similar phenotypic root length responses, however under the −Fe condition the complementing line displayed shorter root than *bts-1* mutant (**Figure3A, Figure S4A**). Measurement of root length under +Fe and –Fe conditions suggest that both the wheat HRZs were able to rescue *the bts-1* insensitivity under both the conditions (**Figure S4B**). In terms of shoot chlorophyll content, the *bts-1* leaves contained lower total chlorophyll content compared to Col-0, Ox-HRZ1, and Ox-HRZ2 under +Fe. As expected and under –Fe, however, the Ox-HRZ1 and Ox-HRZ2 exhibit lower total chlorophyll and Fe content than the *bts-1* mutant (**Figure 3B**). Consistently, ICP-MS analysis showed that the complemented (Ox-HRZ1 and Ox-HRZ2) also accumulated less iron than the *bts-1* mutant under iron-deficient conditions (**Figure 3C**). *bts-1* also exhibited intense Perls staining of the roots compared to the Col-0 and the complemented lines (**Figure 3D, Figure S4C**). Earlier, the enhanced accumulation of Fe in *bts-1* mutant has been attributed to high FCR (Ferric chelate reductase) activity (Hindt et al. 2017). Under +Fe conditions, FCR activity was similar in Ox-HRZ1, Ox-HRZ2, *bts-1* mutant and Col-0. In contrast, under –Fe *bts-1* mutant exhibited significantly higher iron reductase activity (FCR activity) than Col-0 or the complementing lines (**Figure 3E**). At the molecular level, *TaHRZ* genes expressed in *bts-1* could restore the expression of Fe-responsive genes, such as *AtFRO2*, *AtIRT1, AtPYE, AtNAS* and *AtFIT*, to levels similar to Col-0 (**Figure S5**). These results emphasise the conserved role of *TaHRZ* as a possible negative regulator for Fe homeostasis.

**Figure 3:**
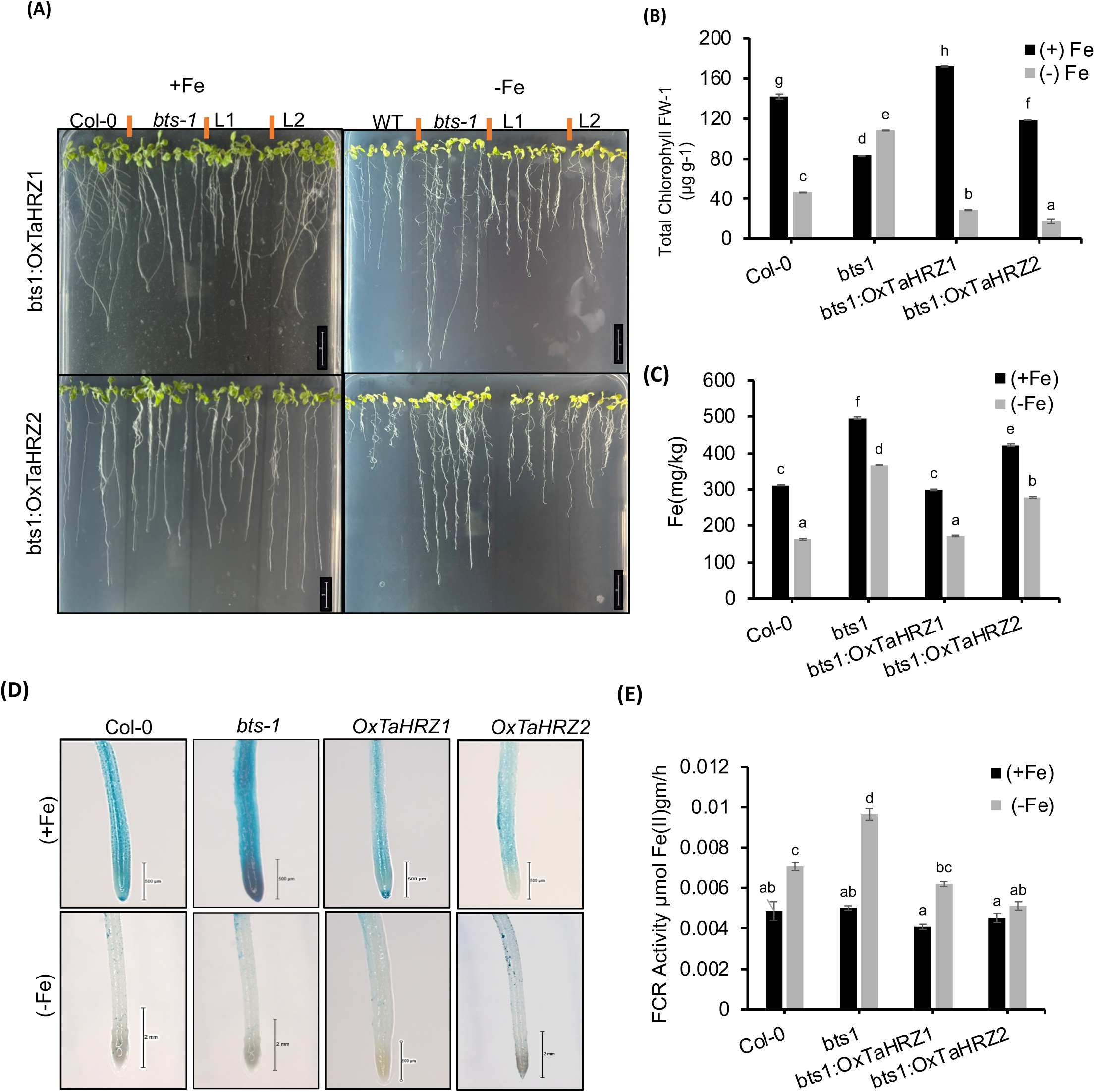
Genetic complementation of *atbts*1 with *TaHRZ*1 and *TaHRZ*2 and its characterization. (A) Phenotypic analysis of *TaHRZ*1 and *TaHRZ*2 complemented lines (Ox-L1 and Ox-L2) in *atbts1* mutant (*bts1*) along with Col-0 (WT). Plants were grown on Hoagland solid medium for 5 days, then transferred to −Fe (200 µM Ferrozine) media for 1 weeks. Under iron deficiency *bts1* mutant exhibits long root growth compared to Col-0. *Arabidopsis thaliana* complementing line *TaHRZ1/HRZ2* reversed the mutant phenotype, resulting in reduced primary root length growth in iron deficiency. (B) Estimation of the total chlorophyll content in the leaves of the treated plants. Chlorophyll assessment was performed using 100mg of leaf tissue. The *atbts1* mutant shown markedly significant increase in the chlorophyll contents under −Fe, +Fe, compared to Col-0 and complementing lines. (C) Total Fe was measured in the seedlings using ICP-MS analysis. (D) Representative images of Fe distribution in Col-0, *bts1*, ox-L1 (HRZ1) and ox-L2 (HRZ2) and *TaHRZ* complementaion *Arabidopsis thaliana* lines, detected by Perls staining. Under −Fe, +Fe, – condition Col-0, *TaHRZ* complementing lines showed less iron accumulation as compared to *atbts1*. (E) Fe chelate reductase (FCR) activity in roots of Col-0, *bts1*, and *TaHRZ1*- and *TaHRZ2*-complemented Arabidopsis lines (ox-L1 and ox-L2) (n=30) in the lines. Letters above the bars indicate significant differences (P < 0.05) between genotypes, as determined by a two-way ANOVA test.

BTS interacts with multiple bHLH IVc proteins to modulate their stability (Li et al., 2021; Long et al., 2010; Selote, et al., 2015). Therefore, we examined whether wheat HRZ proteins exhibit a conserved ability to interact with Fe responsive TabHLH IVc proteins that had previously been identified from wheat (Kumar et al. 2022). Constructs encoding for full-length TaHRZs and their truncated versions (t-HRZs) were transformed in yeast and their protein expression was confirmed by Western blot analysis (**Figure S6A)**. We then assessed individual interactions between TaHRZ and multiple TabHLH proteins, including TaFIT. Both the *TaHRZ1* were able to interact with all the TabHLH and TaFIT homologs from wheat as indicated by growth on the drop-out media -AHLT(x-Gal) (**Figure 4A**). Similary, TaHRZ2 also showed interaction with TaBHLH and TaFIT proteins (**Figure S6B**). BiFC assay further confirmed the strong interaction between TaHRZ1 and TabHLH IVc members (**Figure 4B**). This suggests that TaHRZ encoded proteins possibly possess similar, conserved functional domains that facilitate these protein-protein interactions. Further, we tested these interactions in the absence of the RING and other Zn-finger domains, by performing yeast two-hybrid with the deleted version of the TaHRZ (tHRZ1/2). These truncated versions showed no interaction with TaFIT or any other TabHLH *IVc* TFs (**Figure 4C, Figure S6C**). Our results support the prior observation of the importance of the RING and other Zn-finger domains for protein interaction (Kobayashi, et al., 2013) . Collectively, these results indicate conserved interaction patterns of plant HRZ with bHLH proteins .

**Figure 4:**
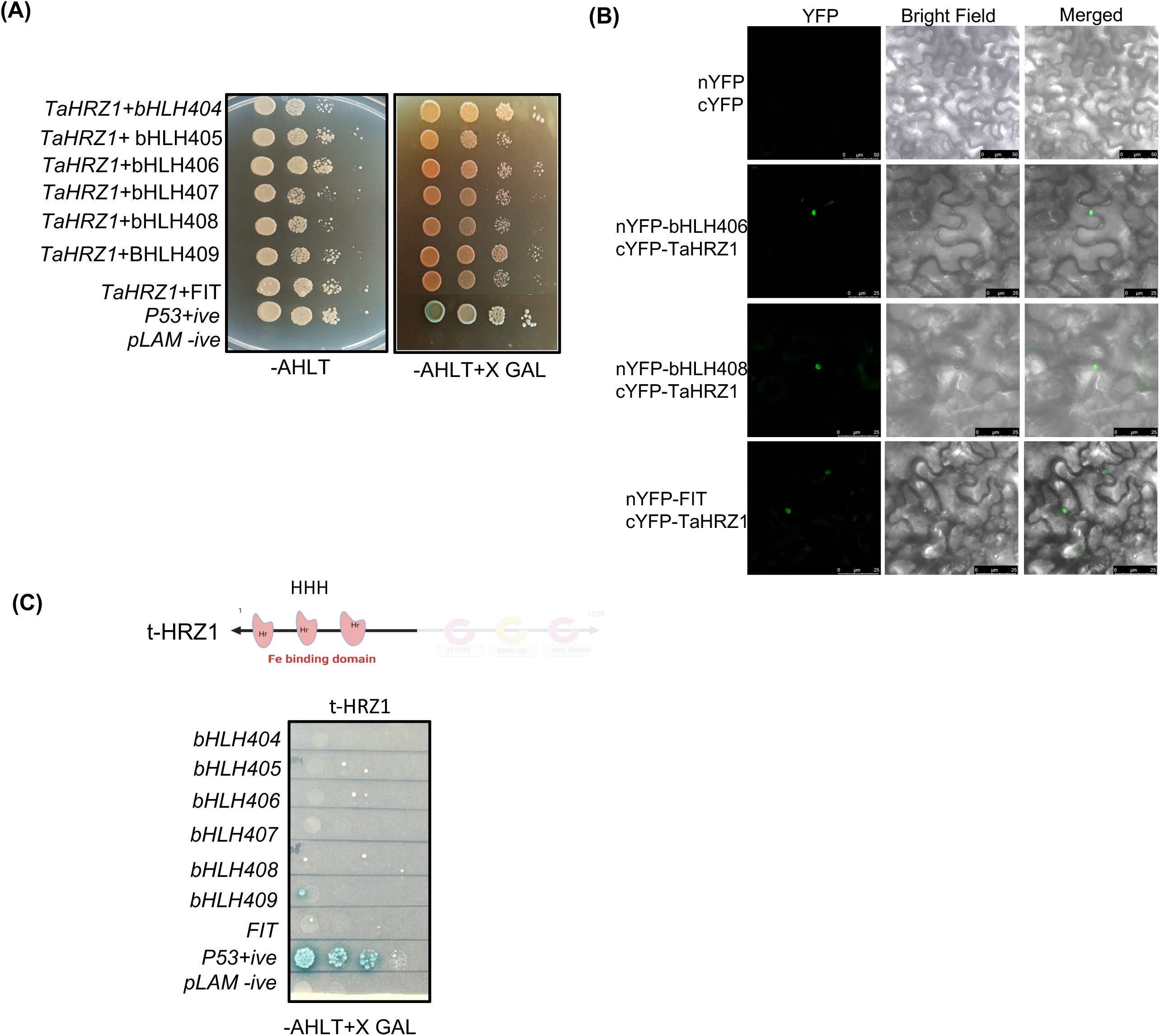
Yeast two-hybrid interactions of TaHRZ1 with the target proteins. (A) Spot growth assay of Y2H GOLD strains expressing a combination of BD- *TaHRZ*1 with AD-BHLH, *TaFIT*. Positive and negative control interaction were also included. (B) Bimolecular fluorescence complementation (BiFC) assay in tobacco leaf cells testing protein–protein interaction of TaHRZ1 with bHLH406, bHLH408 and FIT. The bHLHs were fused to the N-terminal half while TaHRZ1 was fused to the C-terminal half of YFP. Absence of fluorescence in the negative control (upper panel) and the reconstitution of yellow fluorescence shows positive interaction in the nucleus. (C) Interaction of truncated HRZ1 protein (t-HRZ1) with TabHLH IVc (proteins) and TaFIT. Dilutions of OD_600_= 0.100, 0.010, 0.001 and 0.0001 were spotted on solid SD medium or solid SD medium with -LTHA. Plates were incubated at 30 °C for 3 days to assess growth and reporter activity

### GRF4-GIF1 chimeric protein assisted ***g***enome editing of the HHE domain

The HHE domains in the N-terminal region of BTS are essential for protein stability (Pullin et al., 2025; Selote, et al., 2015). Disruption of these domains in *O. sativa* HRZ mutants lead to enhanced Fe in grains (Kobayashi, et al., 2013; Pullin et al., 2025). Therefore, we utilized genome editing to investigate the functional role of the HHE domain encoded by TaHRZ1. To enhance the regeneration efficiencies in Indian wheat, we employed the GRF4-GIF1 chimeric protein for CRISPR-Cas9 editing. We targeted the HHE3 domain of the *TaHRZ1* protein by testing two different guide RNAs (gRNA1 and gRNA2) (**Figure 5A, S7A**). Two individual 20-bp gRNAs were tested, to target the HHE3 domain; however, only one (gRNA2) targeted at exon 7 showed strong in vitro cleavage activity (**Figure S7B)**. This gRNA2 was cloned into the JD633 binary vector system and was driven by the *TaU6* promoter (**Figure 5B**). The resulting construct was introduced into wheat embryogenic callus tissue via *Agrobacterium-*mediated transformation (**Figure S8A,B**). To test the efficiency of the (JD633) GRF4-GIF1 module, we assessed the percentage of callus regeneration for various wheat cultivars (cv.) i.e. Fielder and C306 (**Figure S8C-J and Figure 5C)**. Sixteen independent lines (T_0_) were generated and screened for the presence of the transgene using primer-specific PCR amplification including 9 lines derived from cv. Fielder and 7 lines from cv. C306. (**Table S2 and Figure S9A,B**). The GRF4-GIF1 chimeric protein resulted in enhanced regeneration efficiencies of up to ∼8. in cv. Fielder and 6.4% in cv. C306 (**Table S2**). We conclude that the expression of GRF4-GIF1 module significantly improved regeneration in the Indian wheat cv. C306 and facilitated the efficient recovery of the putative edited events.

**Figure 5:**
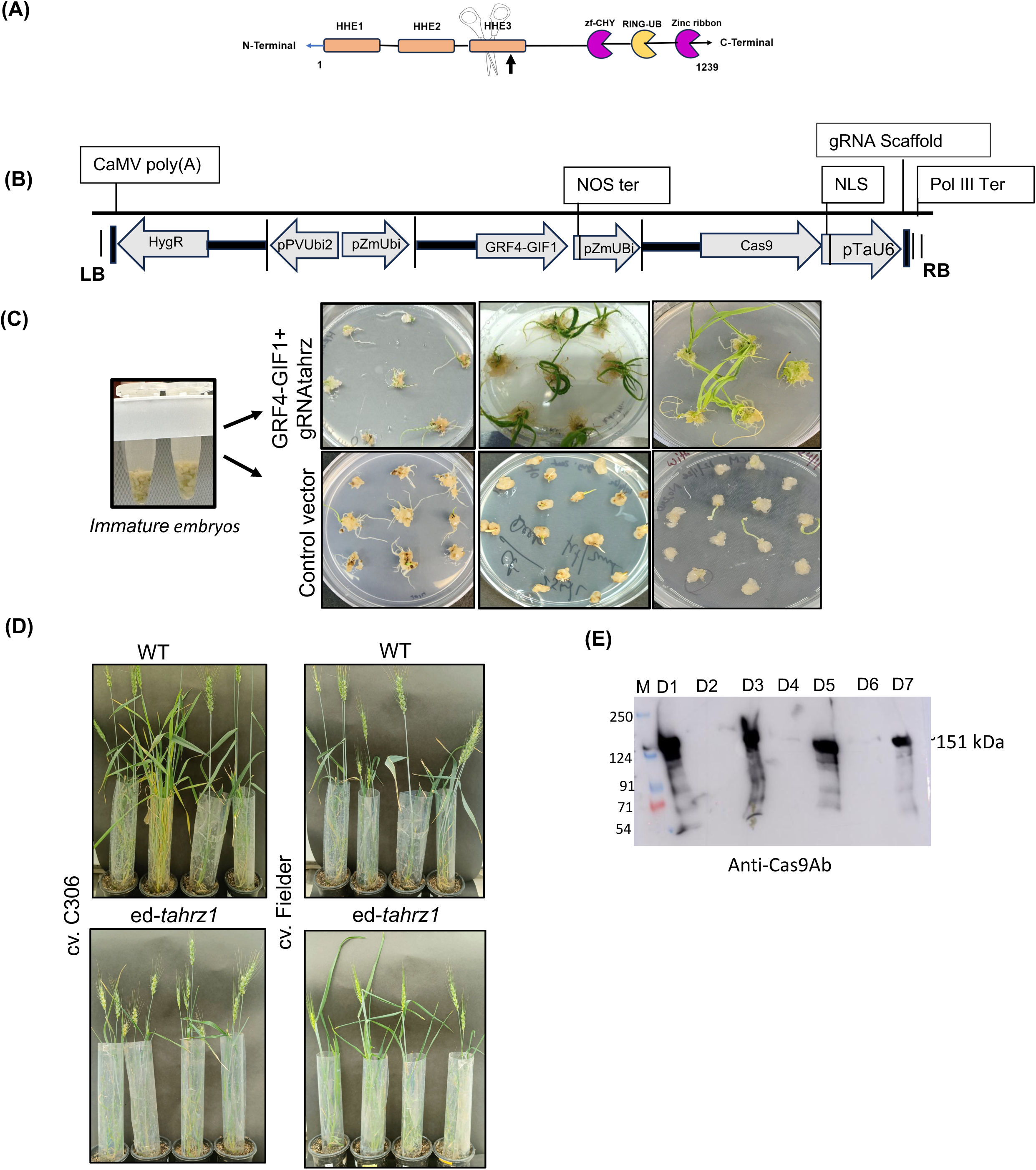
CRISPR-Cas9 mediating editing of HHE3 in *TaHRZ1*. (A) CRISPR-CAS9 editing of HHE3 domain using GRF4-GIF1 to enhance wheat regeneration. HHE3 domain of *TaHRZ*1 was targeted by gRNA for genome editing. (B) The structure that carries the wheat GRF4-GIF1 chimeric protein was derived from JD633 (Addgene). The maize Ubi promoter drives the Cas9 expression, and the gRNA expression is driven by the TaU6 promoter. (C) comparison of post-transformation tissue culture stage in wheat using different constructs. Regeneration response compared to the empty vector (pRGEB32) and GRF4-GIF1 derived clone, across the tissue culture stages. (D) Phenotype of *ed-hrz1* in cv. C306 and cv. Fielder. The pictures were taken post-45 days of growth at the T_1_ generation stage. (E) Western blot analysis to confirm Cas9:TaHRZ1 protein expression in transgenic lines. Total protein was extracted from multiple transgenic lines and probed with anti-Cas9 as the primary antibody and anti-mouse as the secondary antibody. Three lines displayed distinct positive bands at ∼151 kDa, confirming the successful expression of the Cas9:TaHRZ1 fusion protein. Phenotype of *ed-hrz1* (edited lines) in cvs. C306 and Fielder. The pictures were taken post 45 days of growth at T_1_ generation stage.

Next, putative edited lines were identified using the T7 Endonuclease*1* assay, performed on amplicons generated from leaf genomic DNA with primers flanking approximately 150 bp on either side of the gRNA target site (**Figure S9C**). Several putative edited lines from cv. Fielder (D1, D3 and D7) and C306 (C1, C3); and were used for detailed characterization. Putative edited events from the T_1_ plants exhibited normal morphology and development, with no obvious growth penalties associated with editing or transgene expression (**Figure 5D**), indicating successful recovery and establishment of transgenic lines. Western blot analysis of protein extracts from T_1_ leaf tissue confirms the presence of Cas9 in the four transgenic lines of cv. Fielder (**Figure 5E**).

For initial characterization, multiple amplicons derived from the shoot genomic DNA of independent transgenics were generated and subjected to Sanger sequencing. Sanger chromatograms from CRISPR-edited wheat lines of Fielder (D1, D3, D7) and C306 (C3, C11) were analysed to detect indel mutations at the target site. Sanger sequencing of the HHE3 amplicons for the T_1_ plants revealed indel distribution ranged from 1.0 to 16 % for 1 bp deletion. For the 2 bp deletion, the distribution ranged from 1 to 39 % in both cultivars. A predominance of 7 and 11 bp deletions and/or substitutions (+1 bp) was also observed in a few of these lines. Additionally, a larger deletion of 12, 19, 22, and 23 bp was also observed in a few of the edited lines, indicating a frame shift in *TaHRZ*1 and rendering it nonfunctional. Overall, sequence analysis revealed CRISPR/Cas9-induced insertions and deletions ranging from 1 to 23 bp across the edited *TaHRZ1* alleles, with most mutations causing frameshifts and premature stop codons. Indel frequencies were quantified from sequencing data, and high R² values indicated accurate estimation of indel frequencies in the selected lines. **(Figure 6A)**. Indel contribution in these lines occurred near the PAM cleavage site (**Figure 6B)**. Altogether, the Inference of CRISPR Edits (ICE) confirmed successful genome editing. Next, to check the representation of the edits at the T_1_ plant stage, we subjected individual clones to Sanger sequencing (**Figure S10**). These results confirm the successful editing of TaHRZ1 in two genetically distinct wheat cultivars. Furthermore, our data validate the effectiveness of GRF4-GIF1 in promoting higher regeneration in hexaploid Indian wheat cv.C306, thereby enhancing the efficiency of genome editing in wheat.

**Figure 6:**
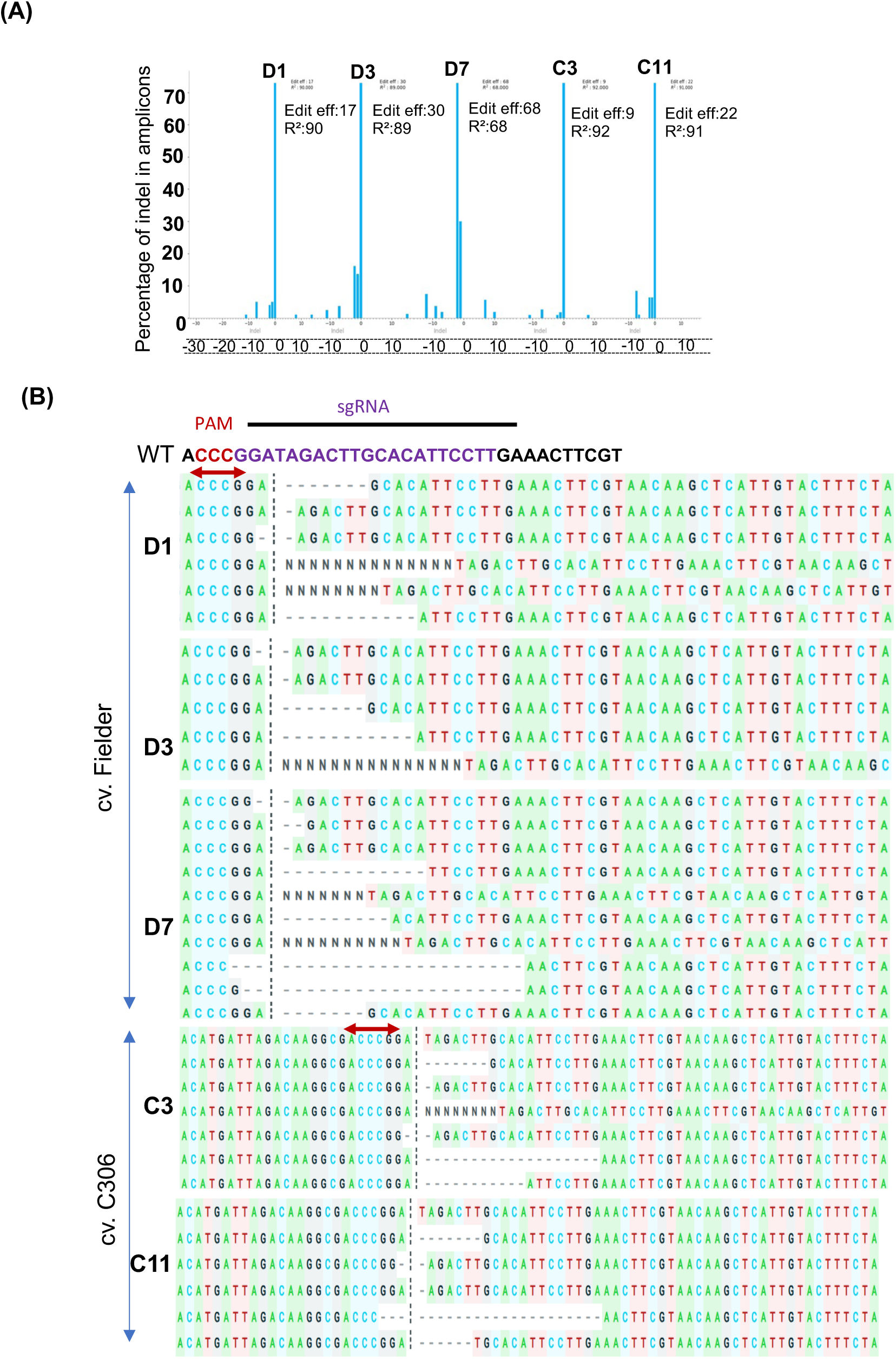
CRISPR edits analysis from Sanger sequencing data of the targeted region near the PAM site to detect interference of CRISPR edits using Synthego. (A) Indel percentages were estimated using Sanger sequencing data, and R² values were used to assess the accuracy of editing. Sanger Chromatograms from CRISPR-edited wheat lines of Fielder (D1, D3, D7) and C306 (C3, C11) were analyzed for indel mutations at the target site. High R² values confirmed efficient and reliable genome editing in the selected lines. Editing efficiency scores of CRISPR-edited wheat lines (Fielder: D1, D3, D7; C306: C3, C11) were calculated based on indel frequencies using Sanger sequencing data. (B) Alignment of the expectations and the different edits containing the insertions and deletions at the target site. R² values represent the confidence in mutation detection and quantification. The red arrow indicates the PAM site (NGG) with the gRNA sequence.

### The defect in TaHRZ1 perturbs Fe distribution in wheat tissue

Previous studies have shown that HRZ proteins regulate grain Fe content in rice grains; however, little is known about their role in controlling Fe spatial distribution within the grain or whether the increased Fe in *TaHRZ1*-edited lines results from enhanced flag leaf remobilization, improved root-to-grain translocation, or both (Kobayashi et al., 2013). The edited lines showed no visible differences in grain morphology compared to controls **(Figure 7A).** To investigate the functional role of *TaHRZ1*, we analysed grain Fe accumulation patterns in *TaHRZ1* edited lines. Perls staining revealed that Fe distribution was notably altered in the edited lines (**Figure 7B, Figure S11A**). In wild-type (WT) plants, Fe was predominantly localised in the aleurone layer and the nucellar projection region of the grain. However, in *TaHRZ1* mutants, Fe accumulation was significantly increased, as reflected by the staining intensity. This may suggest an altered partitioning of Fe during grain filling. ICP-MS analysis showed a 1.52 to 1.96-fold increased grain iron content in edited lines of the two wheat cultivars (**Figure 7C)**. The level of grain phytic acid (PA) ranged from ∼1.2-1.4 % in these transgenic lines **(Figure S11B)**. Interestingly, the molar ratios of Fe:PA were found to be higher for the seeds generated from the transgenic lines when compared to the non-transgenic (**Figure 7D**). Additionally, quantification of other micronutrients showed that Zn and Mn concentrations were moderately increased in the edited lines, whereas Copper (Cu) levels remained unchanged or were slightly reduced (**Figure S12**). These results indicate that *TaHRZ1* editing preferentially affects Fe-associated micronutrient accumulation without causing a broad disruption of metal homeostasis. At the physiological level, while the number of grains per spike remained unchanged, *TaHRZ1* mutants displayed an incremental increase in 1000-grain weight (**Figure 7E,F)**. Despite the altered Fe distribution, no phenotypic differences in overall plant height or shoot architecture were observed between wild-type and *TaHRZ1* mutant lines (**Figure 7G**). These findings imply that *TaHRZ1* disruption improves grain Fe loading without compromising yield-related traits, offering the potential for biofortification strategies to enhance Fe content.

**Figure 7:**
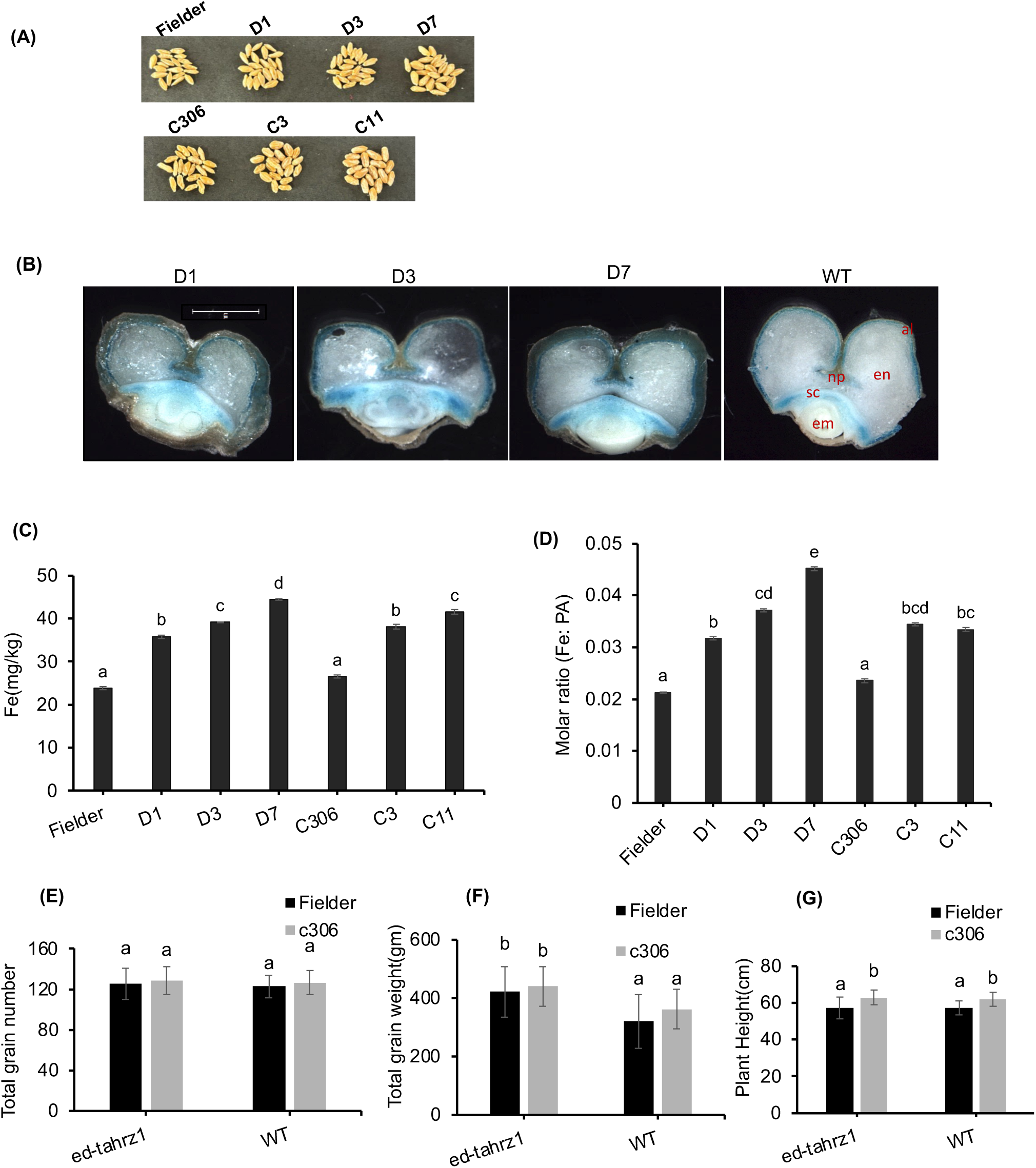
Characterization of wheat CRISPR-Cas9 edits. (A) Phenotype of the grains from the edited lines of two cultivars (B) Representative images of Perl’s Blue staining for transverse cross-sections of transgenic wheat grains (T_1_) at grain maturation. (C) ICP-MS analysis of mature wheat grains (T_1_) (n=4; primary spike). (D) The molar ratios of Iron:Phytate (Fe:PA) were calculated for the mature wheat grains (T_1_) (n=4; primary spike) generated from the transgenic lines and non-transgenic. (E) Average total grain number per plant (n=6). (F) Average total grain weight (1000 grain) in edited lines (GRF4-GIF1). (G) Plant height in cm. Data represent the mean ± SE of six *TaHRZ1* and at least six WT in Fielder, six *TaHRZ1* and six WT in the cv. C306. None of the traits showed any statistically significant differences between the gene-edited and non-transformed control plants. Letters above the bars indicate statistically significant differences (P < 0.05) between genotypes, as determined by one-way ANOVA. (np: nucellar projection; al: aleurone; em: embryo; sc: scutellum; en: endosperm)

### TaHRZ1 influences the expression of Fe homeostasis genes

To further understand the molecular changes associated with *TaHRZ1* disruption, we examined the phenotypic response and expression of key Fe homeostasis regulators in the *TaHRZ1* edited lines. At the phenotypic level, the edited lines of *TaHRZ1 (ed-hrz1)* (2 independent lines for each cv.) show less root growth under +Fe conditions, suggesting that TaHRZ1 could impact root growth (**Figure S13A**). SPAD analysis reveals no significant difference between edited and non-edited seedlings under +Fe and −Fe conditions (**Figure S13B**). At the molecular level, the expression of the *TaHRZ1* was markedly reduced in the edited lines and remained relatively stable under both +Fe and –Fe conditions (**Figure S14A**). Under +Fe conditions, the expression of *TaHRZ2* was not impacted in WT and edited lines of *TaHRZ1*. In contrast, –Fe induces the expression of *TaHRZ2* in the edited lines, but this response was lower when compared to expression in WT seedlings **(Figure S14B)**. Similarly, expression of the Fe Deficiency-Responsive Element Binding Factor 1 (*IDEF1)*, a transcription factor that typically functions upstream in Fe signalling, also shows reduced expression in the edited lines (**Figure S14C**). The suppressed expression of *TaHRZ2* and *TaIDEF1* during –Fe may reflect feedback regulation in response to altered Fe sensing or translocation dynamics caused by the loss of *TaHRZ1* function. Additionally, qRT-PCR analysis revealed a marked upregulation of Fe-related bHLH transcription factor (*TaFIT, TabHLH404, TabHLH407* and *TaIRO3*) in roots and shoots of the edited lines compared to WT plants under Fe deficiency (**Figure 8A-D**). *TaFRO2* and *TaNAS3* did not show significant changes (**Figure 8E,F**). Interestingly, *TaFIT* showed high expression in edited lines even under +Fe conditions. Together, these data suggest that *TaHRZ1* plays a central role in maintaining Fe homeostasis by modulating the expression of both Fe uptake and regulatory genes.

**Figure 8:**
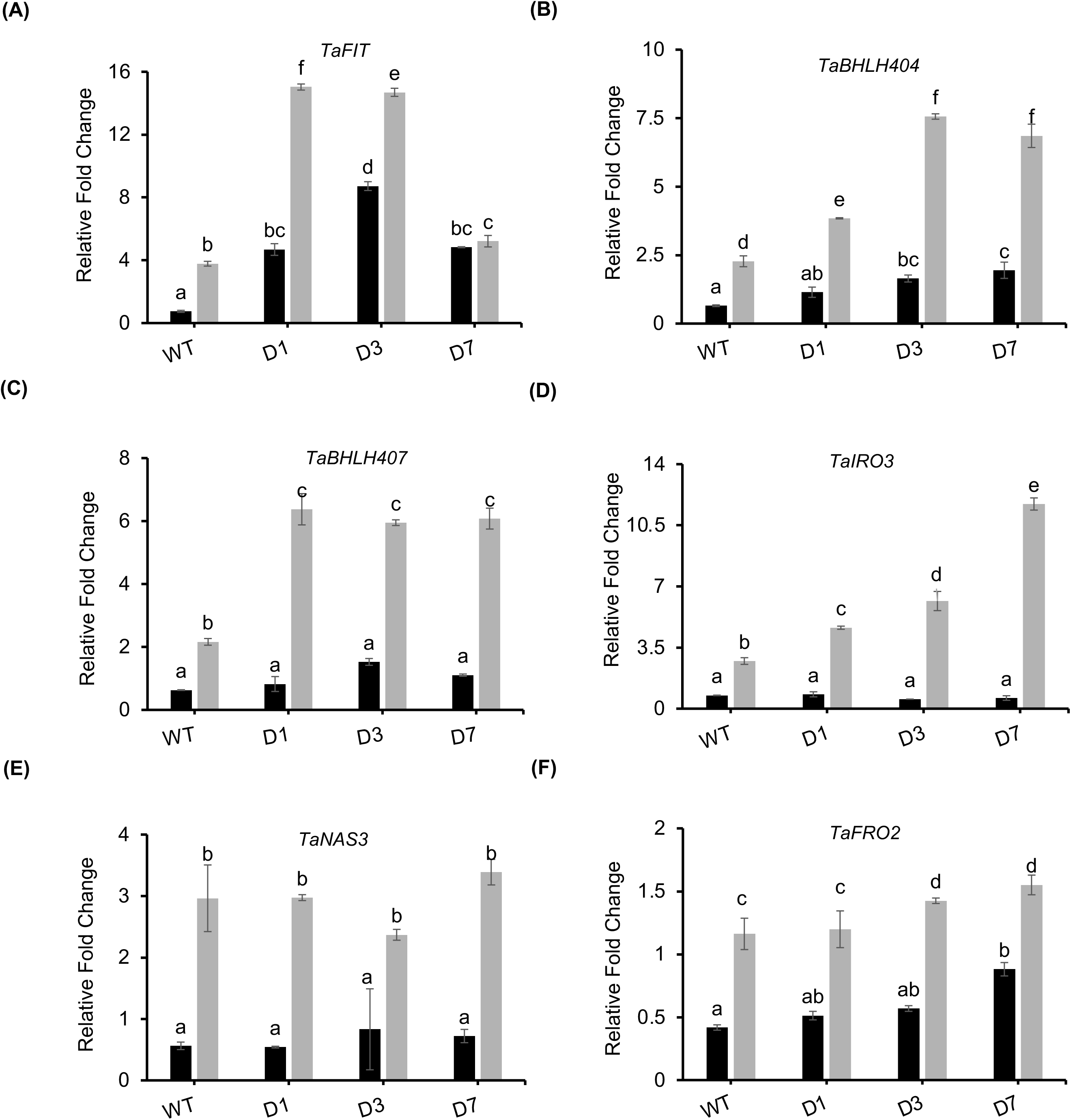
Expression analysis of genes using qRT-PCR in control and edited lines. Relative expression of Fe homeostasis genes (A)*TaFIT*, (B) *TabHLH404*, (C) *TabHLH407*, (D) *TaIRO*3 (E) *TaNAS3* and (F) *TaFRO2* in edited (D1, D3, D7) and control lines (WT) under – Fe and +Fe conditions. Each bar indicates the mean of three replicates with the indicated standard deviation of the mean. C_t_ values were normalized using *TaARF1* as an internal control. Letters above the bars indicate statistically significant differences (P < 0.05) between genotypes, as determined by one-way ANOVA.

## Discussion

Fe is an essential micronutrient required for numerous physiological processes. However, its soil bioavailability is often limited, and maintaining optimal Fe levels within plant tissues requires tight regulation. In graminaceous crops like wheat, Fe homeostasis is critical for plant development and ensuring sufficient micronutrient content in the edible grain parts to meet human nutritional needs. Our investigation into the role of *TaHRZ1*, a wheat homolog of the Fe sensor gene *HRZ*/*BTS*, sheds light on its important regulatory role in Fe distribution and related gene networks in hexaploid wheat. Progress in wheat transformation has lagged behind that of rice and maize, primarily due to the complexity of its genome (Garbus et al. 2015; Wulff & Dhugga 2018). One of the main challenges in wheat gene editing is low regeneration efficiency in current elite cultivars. To overcome this, researchers have employed morphogenetic proteins to enhance embryogenesis and improve regeneration efficiency across several crop species. In particular, the Baby-Boom (BBM) and WUSCHEL (WUS2), WOUND INDUCED DEDIFFERENTIATION1(WIND1) proteins and their combination have significantly boosted transformation frequencies in crops (Laux, Mayer, Berger & Jürgens 1996; Iwase et al. 2017; Lowe et al. 2018). GROWTH REGULATING FACTOR 4 (GRF4) and its cofactor GRF-INTERACTING FACTOR1 (GIF1) have been shown to significantly improve efficiency and speed of regeneration in wheat, triticale, and rice, while also expanding the range of wheat genotype amenable to transformation (Debernardi et al. 2020). In our study, introducing the wheat GRF4-GIF1 module significantly enhanced the regeneration efficiency of calli, resulting in transformation efficiencies ranging from 6.4 % to 8.8% derived from two wheat cvs. C306 and Fielder respectively. GRF4-GIF1 transgenic plants were shown to be fertile, and no obvious developmental defects were observed. In our study, although the transformation efficiency was low, no negative impact was observed for the Indian cv. C306. Further, optimization of the wheat regeneration may be required to adapt and improve the regeneration efficiencies of the transformations. Nonetheless, adapting the GRF4-GIF1 chimeric protein for Indian wheat cv. C306 significantly enhances transformation efficiency compared to previous studies (Aggarwal et al., 2018; Bhati et al., 2016). This reinforces the utility of morphogenetic regulators, such as GRF4-GIF1, for enhancing regeneration efficiency in hexaploid wheat, irrespective of cultivar.

In Arabidopsis and rice, *BTS, BTSL1, BTSL2* and *HRZ* have been characterised as E3 ubiquitin ligases involved in the sensing and degradation of Fe-related transcription factors to maintain homeostasis. While phylogenetic analysis and conserved domain organization indicate that *TaHRZ1* and *TaHRZ2* belong to the HRZ/BTS family, sequence similarity alone is insufficient to confirm equivalent biochemical function in wheat. Recent work by (Pullin et al. 2025) provided biochemical evidence showing that the hemerythrin domain is directly involved in iron binding and sensing, a core feature of HRZ/BTS activity. In combination with our *A. thaliana* complementation assays and protein–protein interaction analyses, these results support a conserved role for *TaHRZ1* in iron sensing and regulatory signalling in wheat. Stable wheat lines expressing tagged *TaHRZ* proteins were not generated due to technical constraints associated with hexaploid wheat transformation. Instead, tissue- and condition-dependent regulation of *TaHRZ1* and *TaHRZ2* was inferred from expression analyses, *A. thaliana bts-1* complementation, and protein–protein interaction studies. Tagged *TaHRZ* lines will be important for future cell-specific localization studies. The wheat *TaHRZ1* fulfils a similar function, albeit with unique regulatory consequences. However, in wheat, the presence of three homoeologs introduces additional layers of complexity, and the high expression of all three in roots points to a root-specific role in Fe sensing and uptake regulation. Interestingly, while *A. bts1* mutants display growth defects under high Fe (Hindt et al. 2017), the wheat *TaHRZ1* mutants have not been explored for this condition yet; future studies may be required in this direction. One of the most interesting aspects of *TaHRZ1* disruption was the coordinated change in the expression of key Fe-regulatory transcription factors. Notably, *TaFIT* and *TaIRO3* were significantly upregulated in the edited lines. *TaFIT* is a central transcription factor in the Fe acquisition pathway (Filiz and Kurt, 2019; Maria N. Hindt et al., 2017; Selote et al., 2015). HRZ proteins act as repressors, so their absence derepresses downstream targets, and this mechanism may result in high expression of known targets such as *TaIRO3* and *TaFIT*. Based on the evidence, it appears that HRZ functions are conserved across dicots and monocots (Li, Watanabe, Gao & Dubos 2023).

Targeted editing of *TaHRZ1* led to striking changes in Fe distribution, with intense Perls staining observed in the scutellum region of the grain. Grain Fe can derive from both flag leaf remobilization and continued root-to-grain translocation; however, as Fe fluxes were not directly traced in this study, we cannot distinguish between these contributions in the TaHRZ1-edited lines. This is particularly notable, as this tissue plays a crucial role in nutrient mobilization during germination. The intense Perls stain in the scutellum suggests altered regulation of long-distance Fe transport, likely due to altered regulatory control over the expression or activity of transporters. Interestingly, this accumulation did not lead to visible growth defects, indicating that *TaHRZ1* is dispensable for vegetative development under normal conditions but is vital for fine-tuning internal Fe movement, especially during seed development. The potential trade-off between enhanced nutrient content and plant fitness or yield is a key concern in biofortification approaches (Ofori, Antoniello, English & Aryee 2022). In our study, *TaHRZ1*-edited lines maintained normal plant height and grain weight, suggesting that the growth processes remained unaffected. These findings highlight *TaHRZ1* as a promising target for improving grain quality without yield penalties, which is a major advantage in crop improvement programs. Our findings also raise the possibility that editing combinations of *TaHRZ1* and *TaHRZ2* may reveal further insights into redundant or complementary roles in Fe regulation. The ability to manipulate Fe distribution within the wheat grain without negatively impacting growth or yield is a significant step forward in biofortification. Current biofortification strategies often rely on overexpression of Fe transporters or chelators, which may result in imbalanced nutrient profiles or impaired growth (Sperotto, Ricachenevsky, de Abreu Waldow & Fett 2012; Pal, Singh & Dhaliwal 2021). Our findings suggest that modulating internal Fe regulators like *TaHRZ1* can enhance Fe accumulation in nutritionally relevant grain tissues. As indicated earlier, the improved Fe:PA molar ratio is indicative of higher bioavailable Fe in the grain (**Figure 7D**). Additionally, changes in the gene expression of *TaFERR1* was observed suggesting that changes in ferritin abundance could potentially influence Fe storage and bioavailability in the edited lines (**Figure S15**). However subsequent studies are required for assessment of the role of HRZ in tissue-specific Fe profiling or employing the isotopic tracer approaches that will be necessary to distinguish between enhanced remobilization and improved root-to-grain translocation in these edited lines.

These efforts, together with further editing strategies (e.g., targeting vacuole Fe transporters or chelators) could potentially redirect Fe to the endosperm to improve its bioavailability further and improve germination under conditions of low Fe availability. Future efforts to gain insight into *TaHRZ* function include determining whether the increased Fe in the scutellum can be mobilized and utilized efficiently during germination. Exploring the combined effects of *TaHRZ1* and *TaHRZ2* double editing could provide deeper insights into redundancy and specificity in the wheat Fe regulatory network. Multiplex gene editing for traits in hexaploids has been routinely used (Zaman, Li, Cheng & Hu 2019). Transcriptomic or proteomic analyses of edited lines could also reveal downstream targets of *TaHRZ1*-mediated regulation, specifically in grains.

Utilizing GRF4-GIF1 to increase transformation efficiency we have shown that *TaHRZ1* is a key regulator of Fe accumulation and Fe distribution in wheat. Moreover, we have shown that disrupting TaHRZ1 enhances Fe accumulation in grain scutellum without compromising yield-related traits. The accompanying changes in *TaFIT*, *TaIRO3*, and *TaIDEF1* expression highlight the central role of *TaHRZ1* in the Fe regulatory network. Further studies required generating wheat grains with desired heritable edits and assessing the double mutants of wheat HRZs to decipher additional and regulatory mechanisms. Altogether, this study provides insights into the molecular mechanisms underlying HRZ-mediated nutrient regulation and establishes genome editing of HRZ as a viable strategy for biofortifying wheat grains. These findings pave the way for developing nutrient-dense wheat varieties, addressing global micronutrient deficiencies and enhancing food security. These results provide a promising foundation for developing nutritionally enriched wheat varieties through precise genome editing strategies targeting endogenous regulators of micronutrient homeostasis.

## Acknowledgements

The authors thank the Executive Director of NABI for facilities and support. The NABI-CORE grant to AKP supported this work. DT is thankful to UGC for Ph.D. fellowship. DBT-eLibrary Consortium (DeLCON) is acknowledged for providing timely support and access to e-resources for this work.

## Contributions

Conceptualisation: DT and AKP; Project administration: AKP; Investigation: DT, DKJ,AK, HB, KA, RJ, TL, VM,; Formal analyses: DT, AKP, DKJ, VM; Visualization: DT, DKJ, PY, SBS, VM; Resources: AKP, KA, AK; Methodology: All authors. Supervision: AKP, AK. Writing- Original draft: AKP, DT; Writing- Review and Editing: AKP, TL, DKJ, PY, DT, SBS, VM, AK; Funding Acquisition: AKP.

## Conflict of interest

The authors declare that they have no known competing financial interests or personal relationships that could have appeared to influence the work reported in this manuscript.

## Notes

### Competing Interest Statement

The authors have declared no competing interest.

### Summary of Updates

New Data has been added in Figure 4 and multiple SUpplementary information as Figures has been submitted now for better clarity and larger readership.

### Funding

- Department of Biotechnology: NABI-CORE

