## Supplementary Figures for "CRISPR/Cas9 editing of the wheat iron sensor *TaHRZ1* confirms its conserved role in iron homeostasis and allocation in grains"

(A)

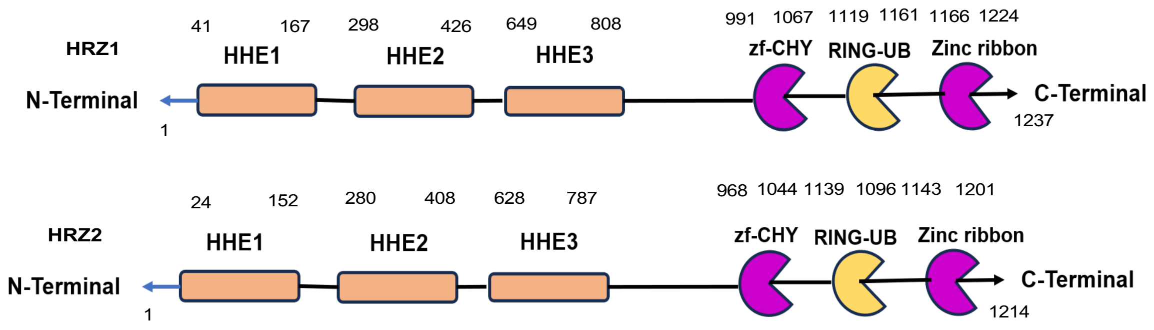


(B)

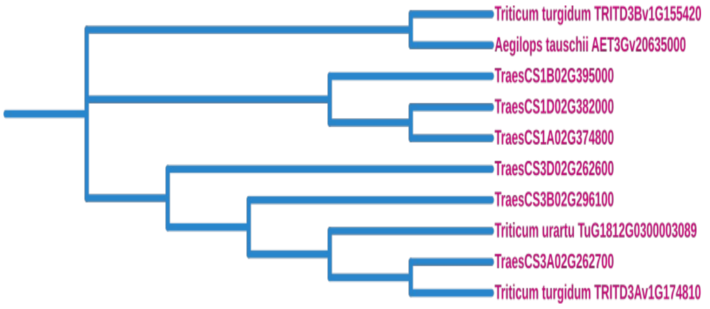


| S. No | Protein Identity matrix | *OsHRZ1* | *AT3G18290* | *TraesCS3B02G296100* | *TraesCS3A02G262700* | *TraesCS3D02G262600* | *OsHRZ2* | *TraesCS1B02G395000* | *TraesCS1A02G374800* | *TraesCS1D02G382000* |
|---|---|---|---|---|---|---|---|---|---|---|
| 1 | *OsHRZ1* | 100 | 58.49 | 88.59 | 88.73 | 88.64 | 76.69 | 73.94 | 73.86 | 73.90 |
| 2 | *AT3G18290* | 58.49 | 100 | 61.16 | 61.72 | 61.56 | 65.71 | 60.7 | 60.39 | 60.08 |
| 3 | *TraesCS3B02G296100* | 88.59 | 61.16 | 100 | 99.09 | 98.68 | 78.03 | 75.27 | 75.58 | 75.6 |
| 4 | *TraesCS3A02G262700* | 88.73 | 61.72 | 99.09 | 100 | 98.79 | 78.79 | 75.59 | 75.9 | 75.83 |
| 5 | *TraesCS3D02G262600* | 88.64 | 61.56 | 98.68 | 98.79 | 100 | 78.42 | 75.18 | 75.57 | 75.51 |
| 6 | *OsHRZ2* | 76.69 | 65.71 | 78.03 | 78.79 | 78.42 | 100 | 81.63 | 81.48 | 81.38 |
| 7 | *TraesCS1B02G395000* | 73.94 | 60.7 | 75.27 | 75.59 | 75.18 | 81.63 | 100 | 96.77 | 96.62 |
| 8 | *TraesCS1A02G374800* | 73.86 | 60.39 | 75.58 | 75.9 | 75.57 | 81.48 | 96.77 | 100 | 98.63 |
| 9 | *TraesCS1D02G382000* | 73.90 | 60.08 | 75.6 | 75.83 | 75.51 | 81.38 | 96.62 | 98.63 | 100 |

**Supplementary Figure S1:** Full-length cDNA sequences and predicted protein structures of *TaHRZ1* and *TaHRZ2*. (A) The full-length cDNAs of *TaHRZ1* and *TaHRZ2* are 3714 bp and 3645 bp, respectively. Their open reading frames encode proteins of 1237 and 1214 amino acids. (B) Phylogenetic analysis and identity matrix of HRZ proteins from diploid, tetraploid, and hexaploid wheat and their progenitor species. HRZ homologs from *Triticum urartu* (A genome), *Aegilops tauschii* (D genome), tetraploid wheat (AABB), and hexaploid bread wheat (*Triticum aestivum*, AABBDD) cluster according to their A, B, and D genomic origins, reflecting the evolutionary conservation and divergence of HRZ genes during wheat polyploidization.

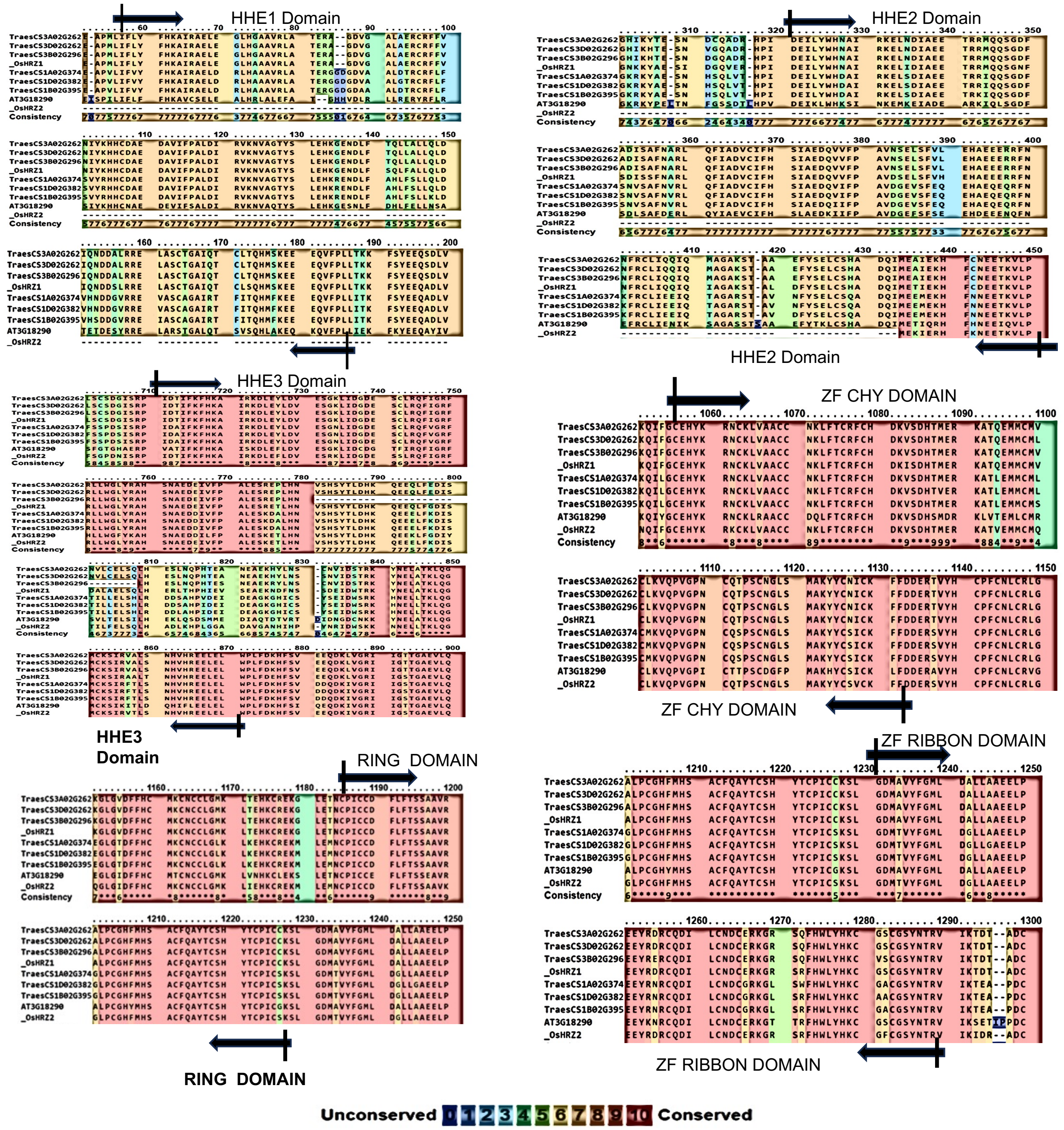


**Supplementary Figure S2** : Multiple sequence alignment of HRZ proteins showing conserved N-terminal HHE domain and C-terminal ZF-CHY/RING-finger (ZF-ribbon) domain in wheat (*TaHRZ*), Arabidopsis (*AtHRZ*), and rice (*OsHRZ*). Conserved residues within the HHE motifs and zinc-coordinating cysteine and histidine residues of the ZF domain are highlighted.

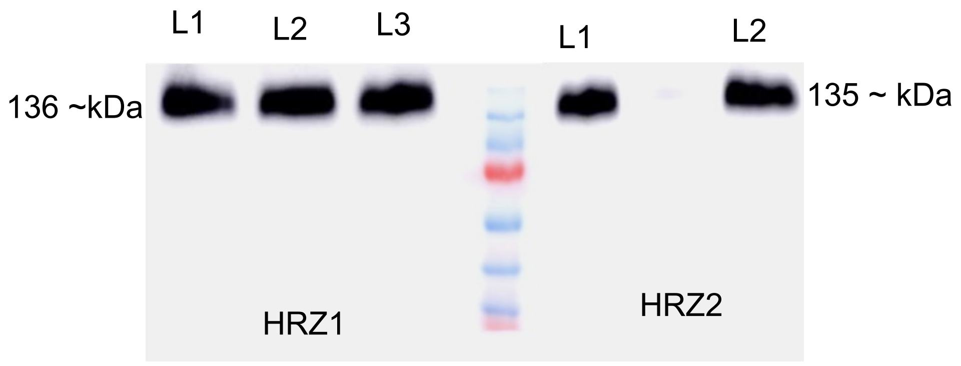


**Supplementary Figure S3 :** Characterization of *atbts* complementation with TaHRZ1/HRZ2 under –Fe (200µM Ferrozine),and control condition (80 µM). (A) Multiple lines were confirmed for protein expression by Western analysis of 6XHis-tagged-HRZ proteins at ‘C terminal.

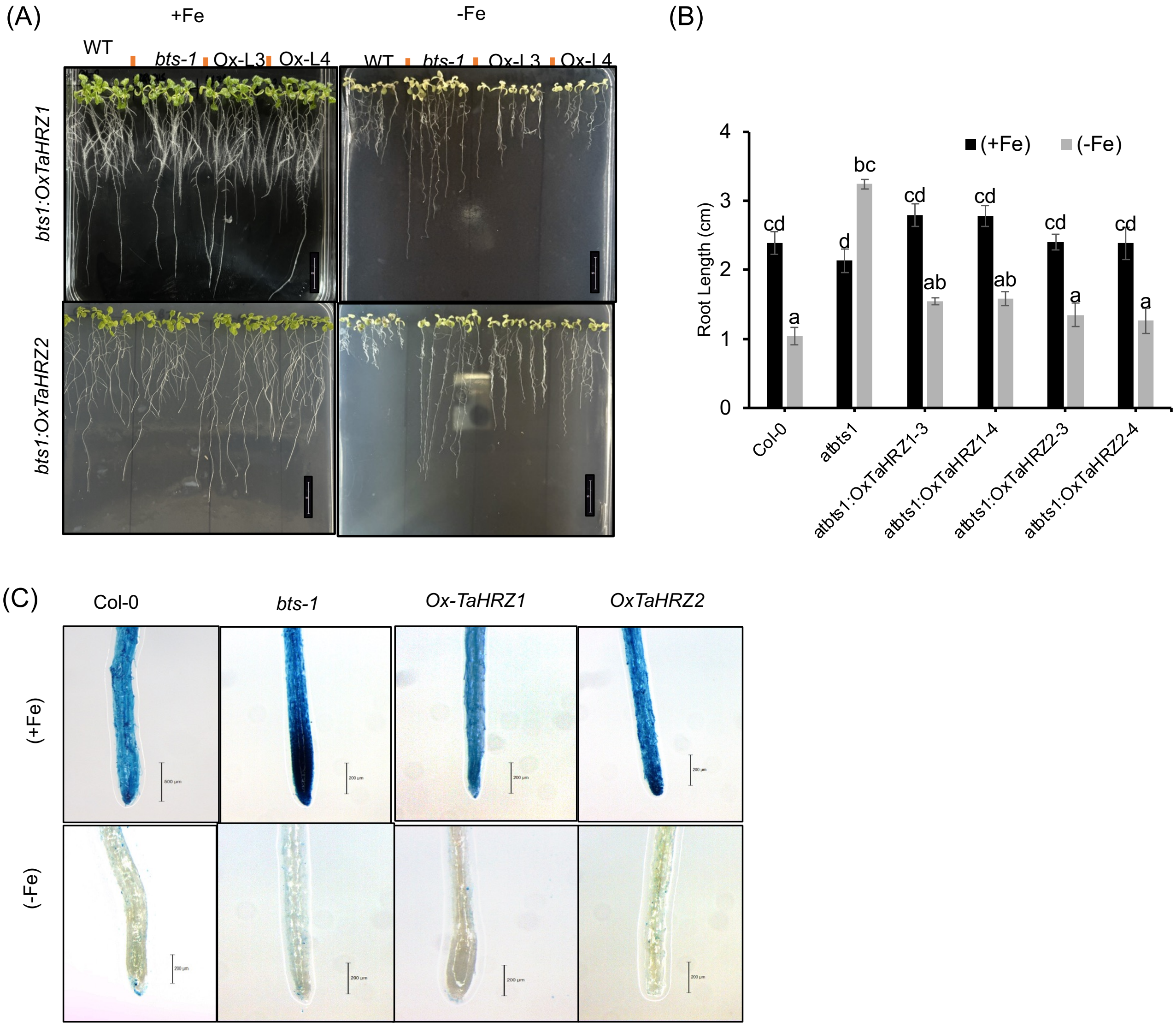


**Supplementary Figure S4**: Genetic complementation and characterization of the atbts1 mutant by *TaHRZ1* and *TaHRZ2*. (A) Phenotypic analysis of *TaHRZ1*- and *TaHRZ2*-complemented Arabidopsis lines (Ox-L3 and Ox-L3, respectively) in the atbts1 (bts1) mutant background, along with Col-0 (WT). (B) root length measurement of the lines. Complementation with *TaHRZ1* or *TaHRZ2* restored the wild-type phenotype, resulting in reduced primary root length under –Fe conditions.(C) Representative Perls staining images show iron distribution in Col-0, bts1, Ox-L3 (*TaHRZ1*), and Ox-L4 (*TaHRZ2*) lines. Under both iron-sufficient (+Fe) and iron-deficient (–Fe) conditions, Col-0 and *TaHRZ*-complemented lines displayed lower iron accumulation compared to the atbts1 mutant. Letters above the bars indicate statistically significant differences among genotypes ($P < 0.05$), as determined by two-way ANOVA.

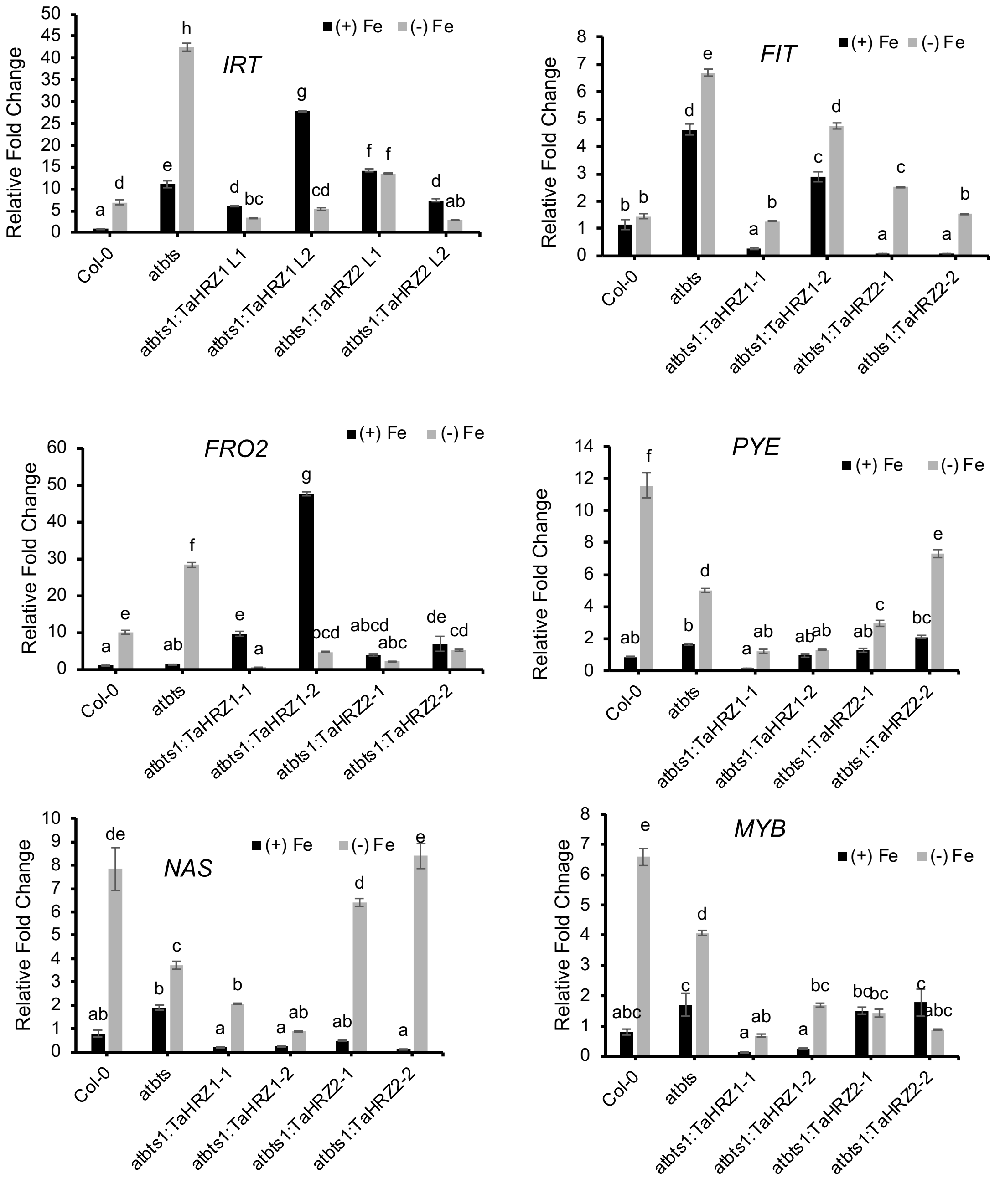


**Supplementary Figure S5:** Expression analysis of known *Arabidopsis* Fe-responsive genes in Col-0, atbts mutant and complemented lines with TaHRZ1/TaHRZ2 under control (+Fe) and iron deficiency (-Fe) conditions.

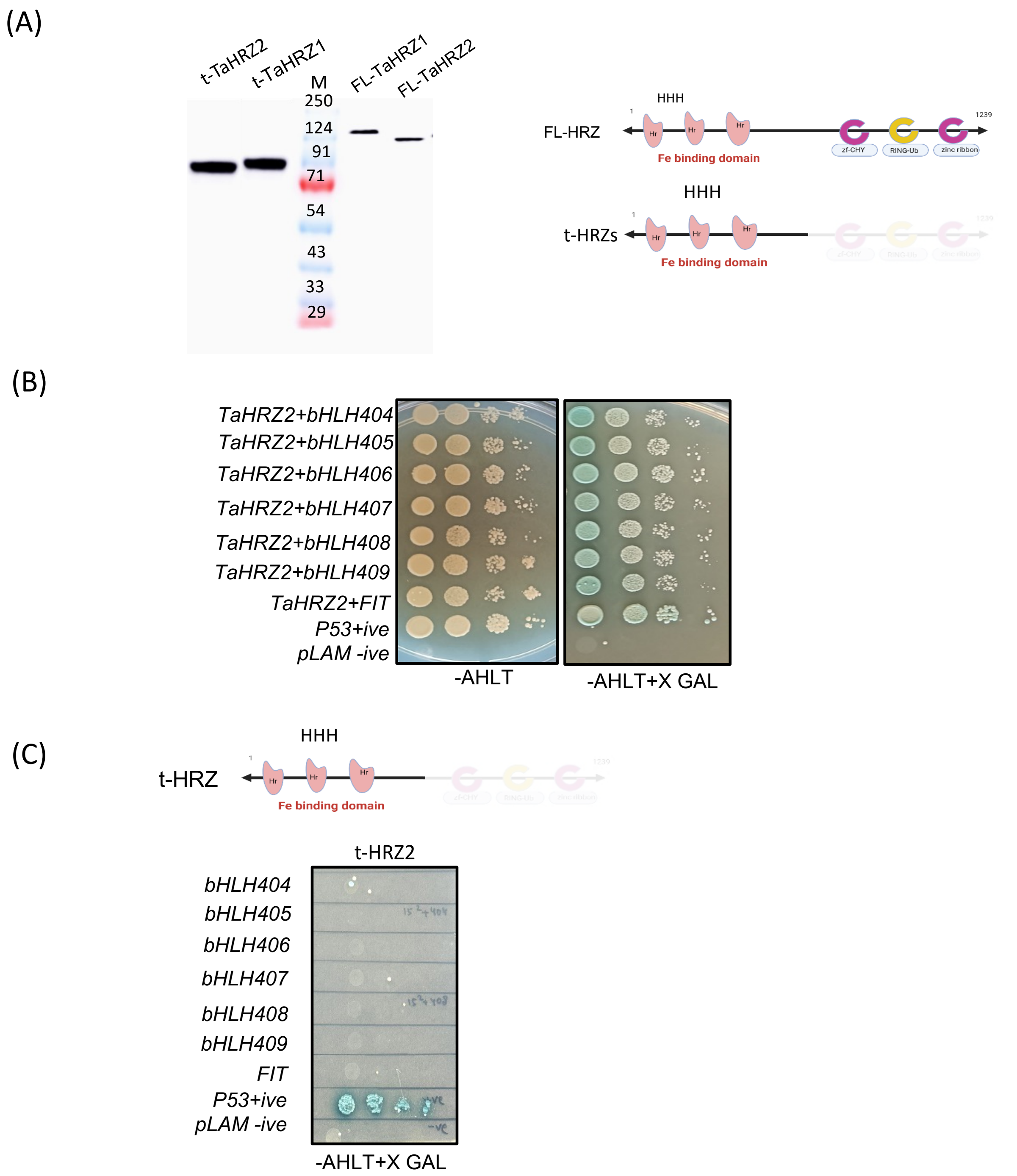


**Supplementary Figure S6:** Yeast two-hybrid interactions of TaHRZ2 with the target proteins**.** (A) Western confirmation of the expression of full length TaHRZ TaHRZ2 proteins (FL-HRZs) and their truncated versions (t-HRZs : without fused C terminal ringh domain) tagged with c-Myc in the yeast strains. (B) Spot growth assay of Y2H GOLD strains expressing a combination of BD-*TaHRZ*2 with AD-*TabHLH*, *TaFIT*. Positive and negative control interaction were also included. (C) Sport assays for the t-HRZ2 interaction with multiple bHLH interating partners. Dilutions of $OD_{600}$= 0.100, 0.010, 0.001 and 0.0001 were spotted on solid SD medium or solid SD medium with -LTHA. Plates were incubated at 30 °C for 3 days to assess growth and reporter activity

(A)

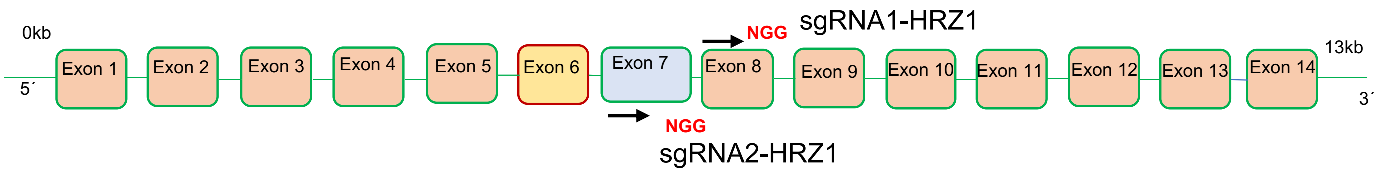


(B)

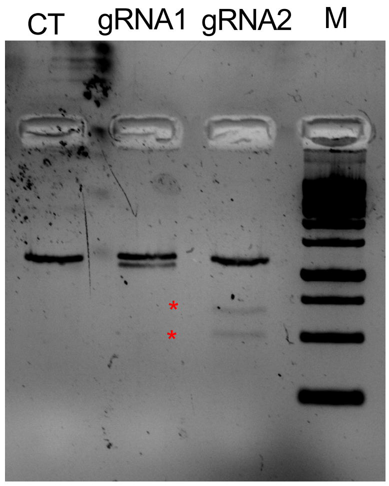


HRZ1_gRNA1: 1318= (771+627)
HRZ1_gRNA2: 1287=(500+687)

**Supplementary Figure S7:** (A) CRISPR-Cas9 target sites within the *TaHRZ1*, with PAM (NGG) sequences located in exon 7 and exon 8 selected for editing. (B) In vitro Cas9 cleavage analysis with genomic fragment for regions targeting wheat HRZ1 and HRZ2 genes at the conserved regions of A,B&D chromosomes. Lane 3 (marker), Uncut amplicon for for target : CT was PCR-amplified and digested with Cas9+gRNAtarget1 and Cas9+gRNAtarget2, respectively. Stars denote digested bands.

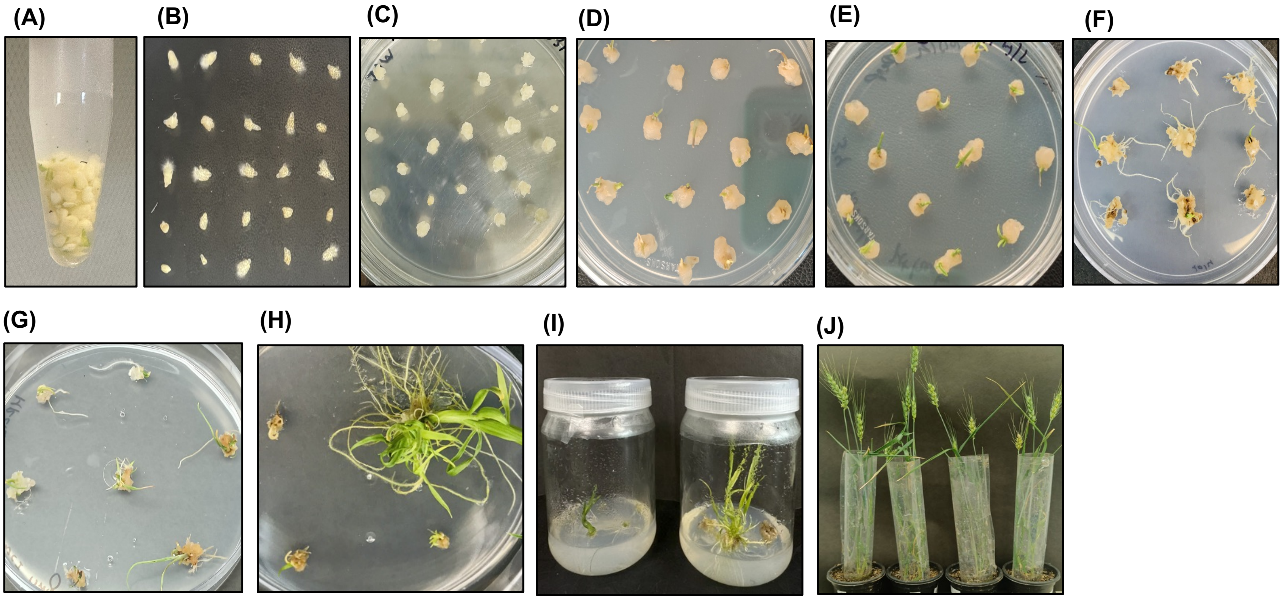

**Supplementary Figure S8:** Multiple stage of agrobacterium mediated transformation for control vectors or vector carrying GRF4-GIF1. (A) Immature embryo harvested from 14-16 DAA(B) Embryo infected with Agrobacterium strain AGL1 (C) Resting on co- cultivation for 3days (D) Infected calli on callus induction media (E) and (F) Control vector and GRF4-GIF1 infected calli on selection media (15 mg/lit) after 1 cycle of subculturing (G) and (H) Control vector and GRF4-GIF1 infected calli on selection media(30mg/lit) after 2 cycle of subculturing (I) Seedling selected on selection media kept for rooting regeneration media (J) followed by harvesting on vermiculite.

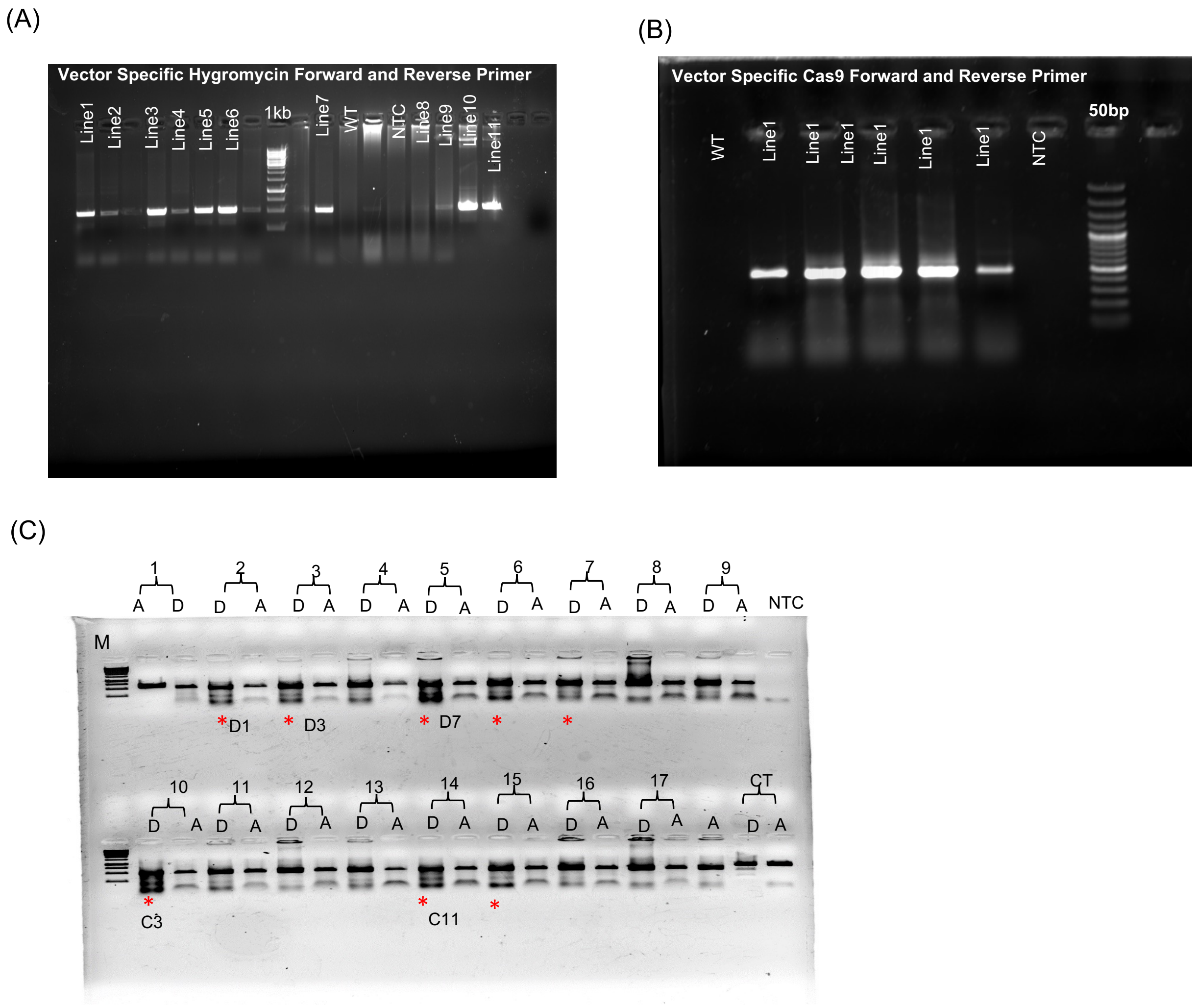


**Supplementary Figure S9:** Molecular screening of CRISPR Cas9 generated TaHRZ1 transgenic lines. Gel images showing the PCR screening of TaHRZ1 transgenic lines. Two sets of primers were used for amplification: (A) The first set targeting the vector-specific hygromycin forward and reverse primers (approximately 500 bp), (B) the second set using vector-specific cas9F and gene-specific reverse primers (approximately 300 bp). Wild-type (WT) genomic DNA was used as a negative control. (C) T7Endonuclease1 (T7E1)assay for screening of putative mutation in the targeted region of the amplicons. Red * indicates the sample those were processed further for analysis because of the banding pattern.

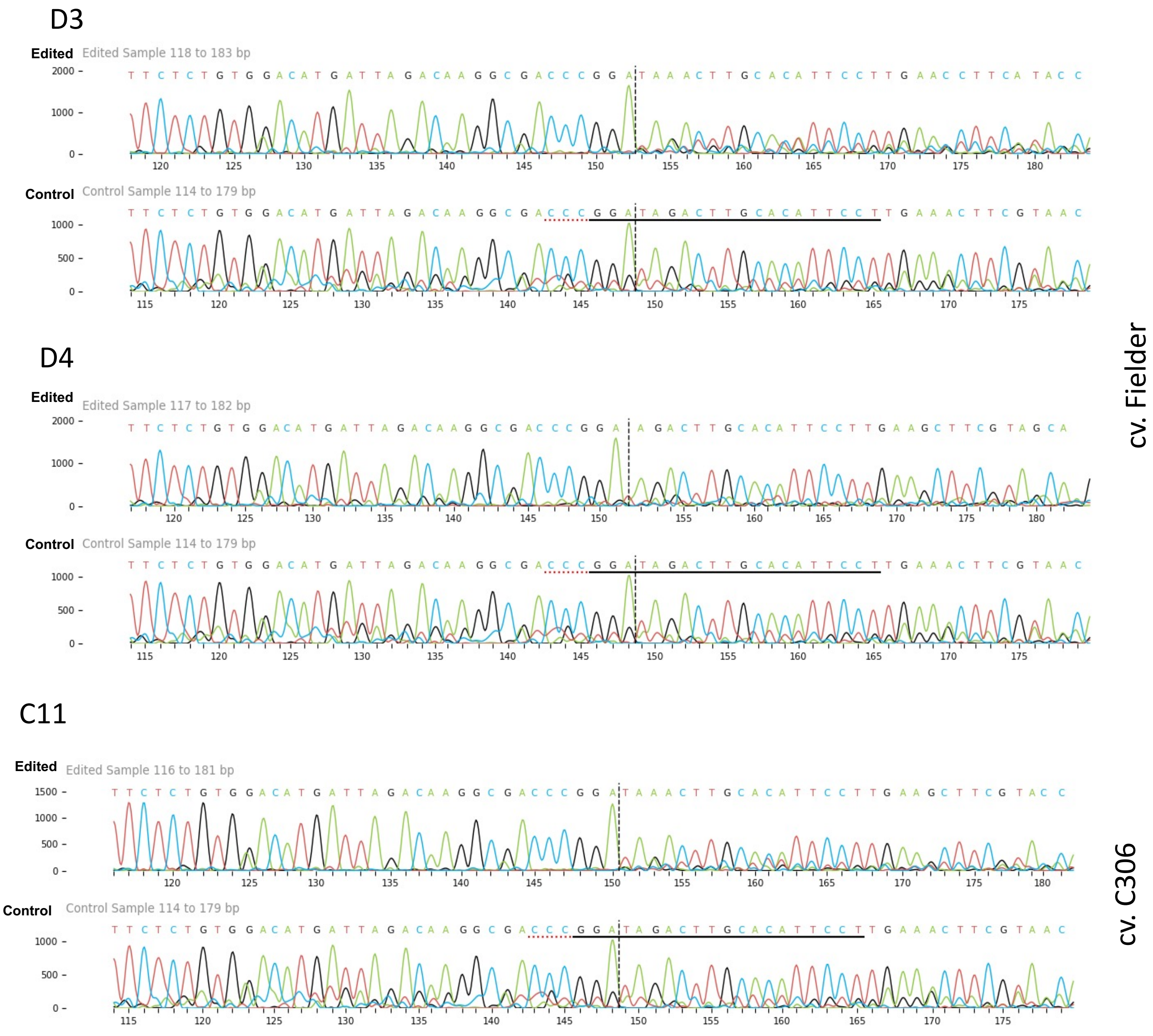


**Supplementary Figure S10:** Chromatogram alignment of the sequence PCR amplicon from the *TaHRZ1* transformants.

**(A)**

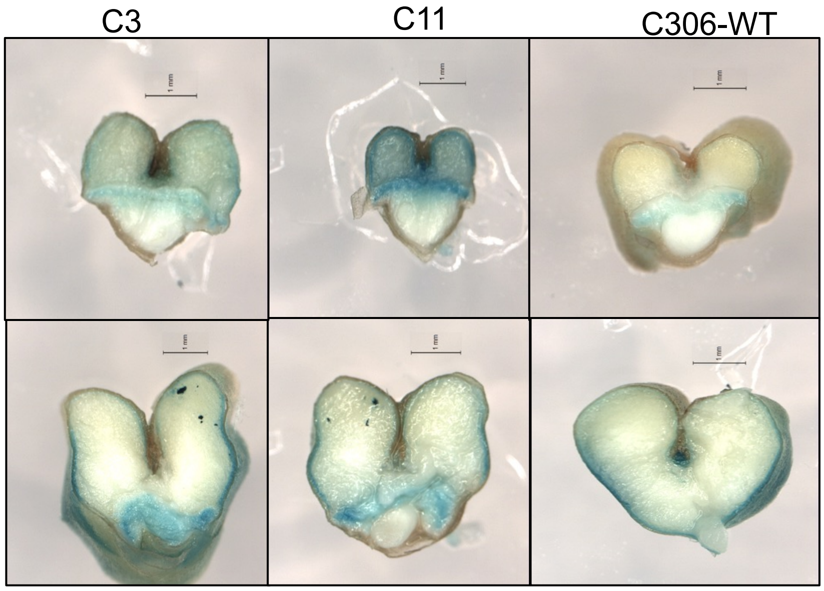


**(B)**

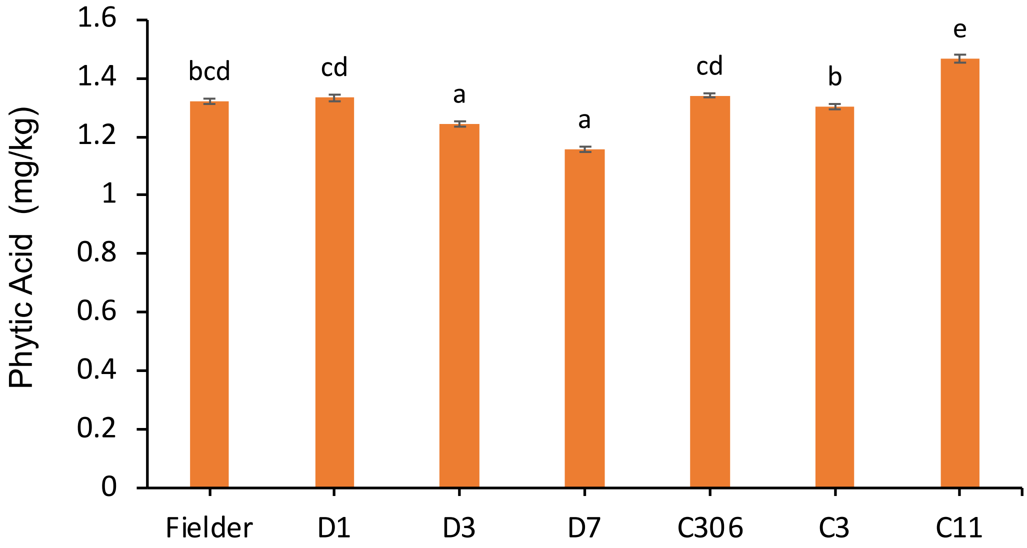


**Supplementary Figure S11:** Iron staining and phytic acid measurements in wheat grains (A) Perls staining to visualize iron localization in grains of *TaHRZ1* transformants in C306 cultivar compared to their respective controls.(B) Estimation of grain phytic acid (PA) content in transgenic and non-transgenic wheat lines. Grain phytic acid (PA) levels were quantified from mature seeds of CRISPR-edited lines (Fielder: D1, D3, D7; C306: C3, C11) and compared with their respective non-transgenic controls. Data represent mean ± SD (n = 3). Statistical significance was evaluated using one way ANOVA.

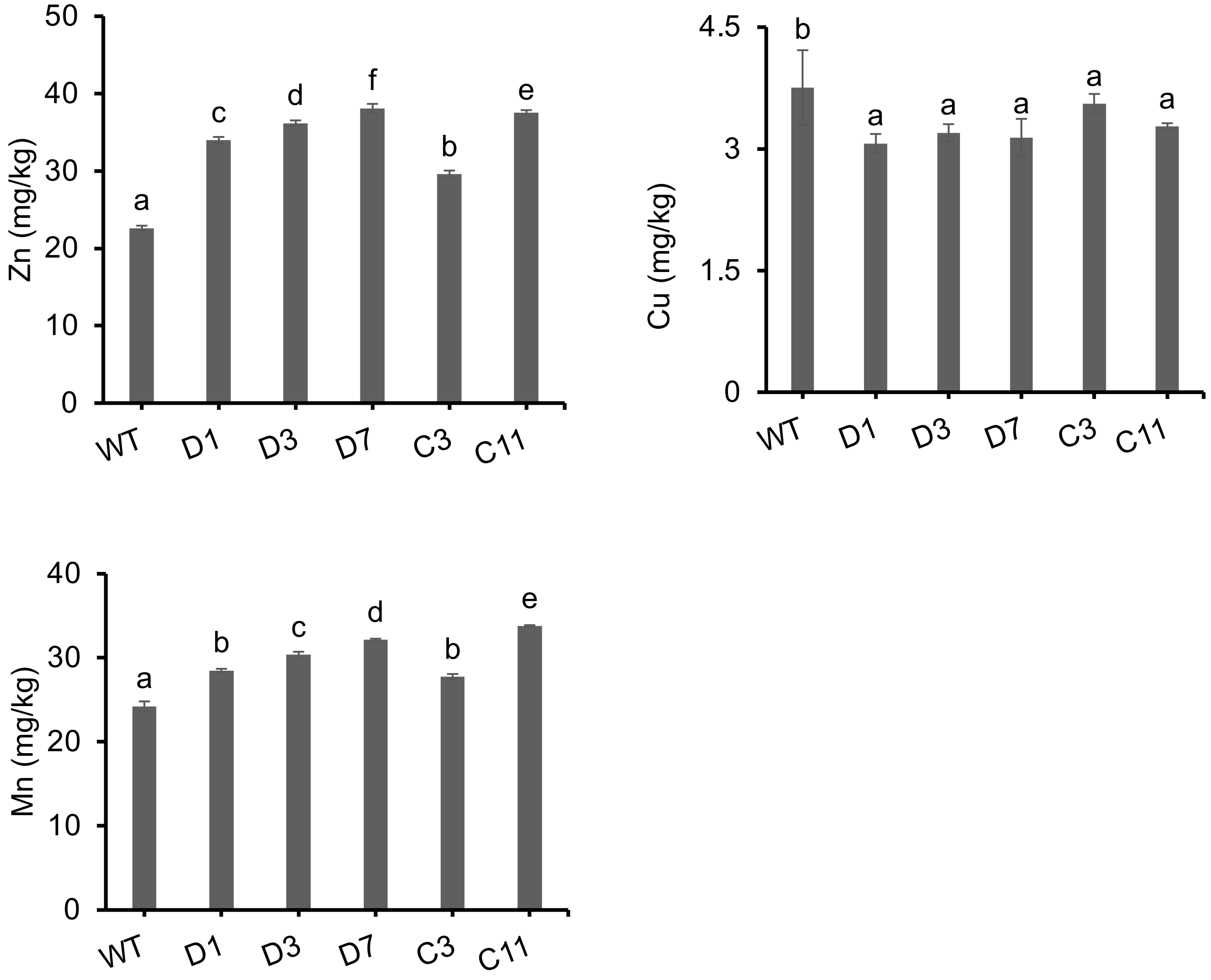


**Supplementary Figure S12**: ICP-MS analysis of Zn, Mn, and Cu concentrations in mature wheat grains (T1 generation, primary spike; n = 4). Letters above the bars indicate statistically significant differences (P < 0.05) between genotypes, as determined by one-way ANOVA.

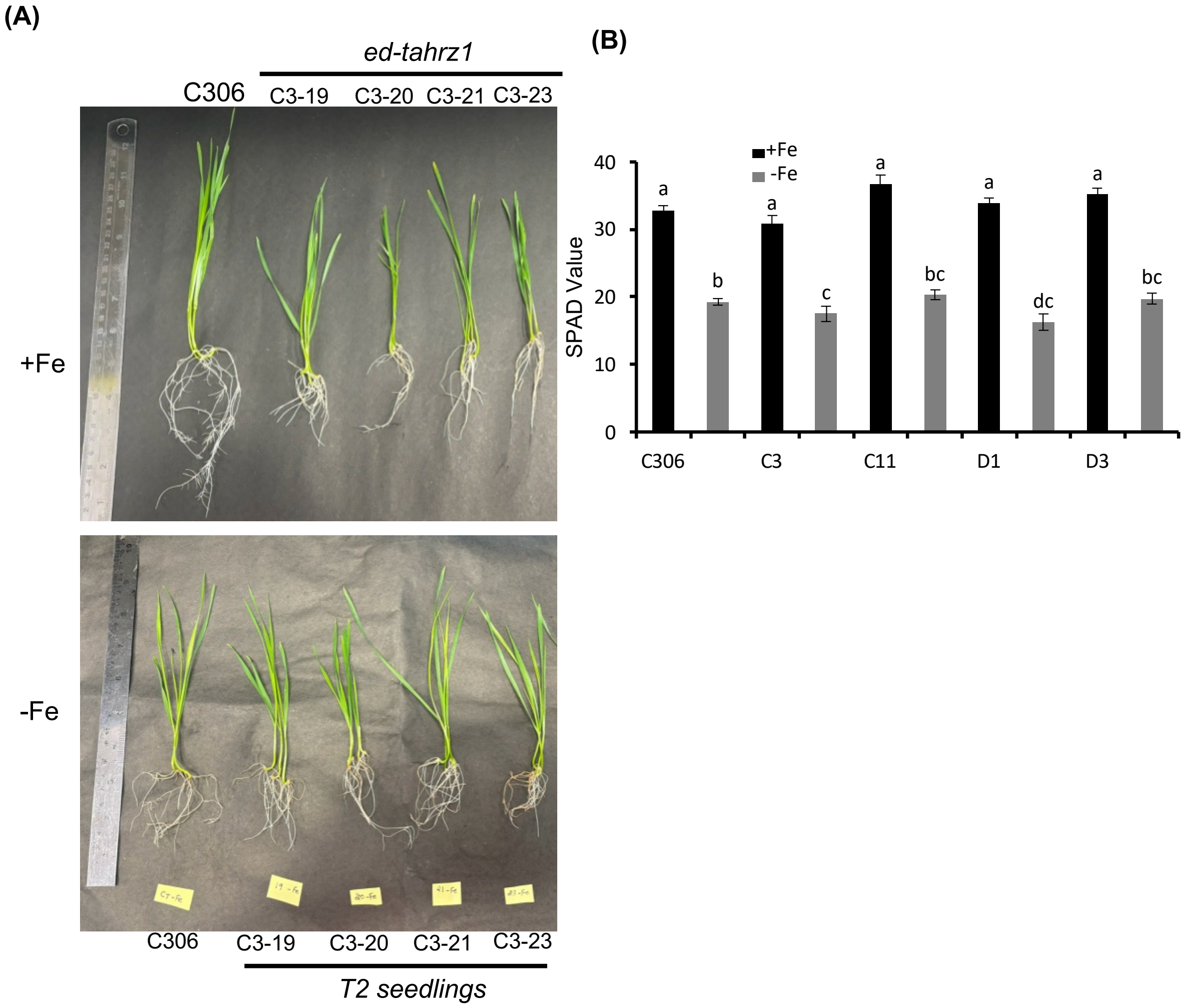


**Supplementary Figure S13:** (A) Phenotype of Control (cv. C306) and the edited lines (T2) of wheat ed-tahrz1 under control (+Fe) and during Fe-deficiency condition (-Fe) after 8 days of experiments (n=6). (B**)** SPAD values were taken on the young leaves of the seedlings (n=4) and 3 different areas of leaves. Statistical significance was evaluated using one-way ANOVA.

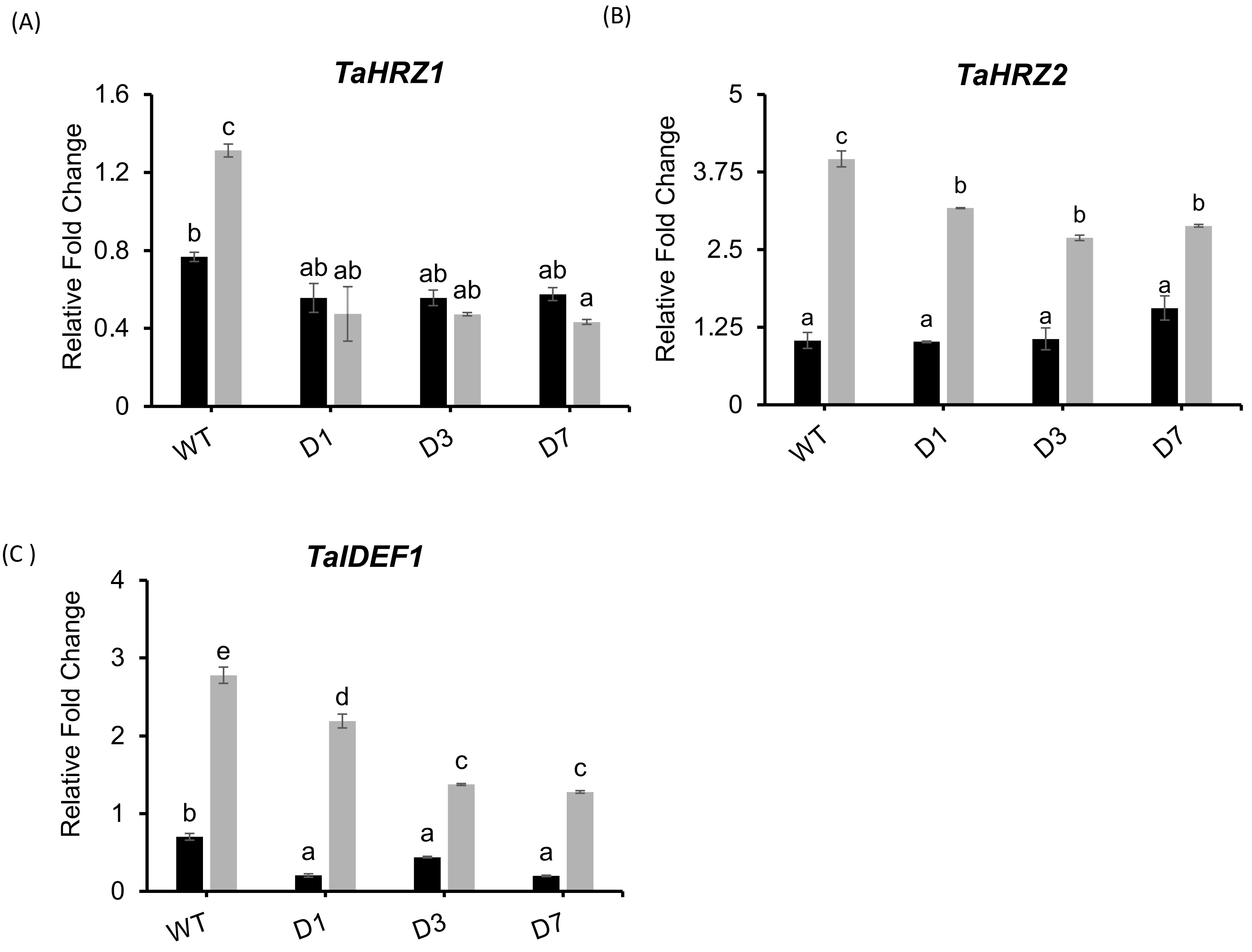


**Supplementary Figure S14**: qRT-PCR–based expression analysis of Fe homeostasis–related genes in WT and TaHRZ1-edited seedlings. Relative transcript levels of (A) *TaHRZ1*, (B) *TaHRZ2*, and (C) *TaNAS3* were analyzed in edited lines (D1, D3, D7) and WT plants grown under Fe-sufficient (+Fe) and Fe-deficient (–Fe) conditions. Each bar represents the mean ± SD of three biological replicates. Gene expression levels were normalized to *TaARF1* as an internal reference. Different letters above the bars indicate statistically significant differences ($P < 0.05$), as determined by two-way (two-factor) ANOVA

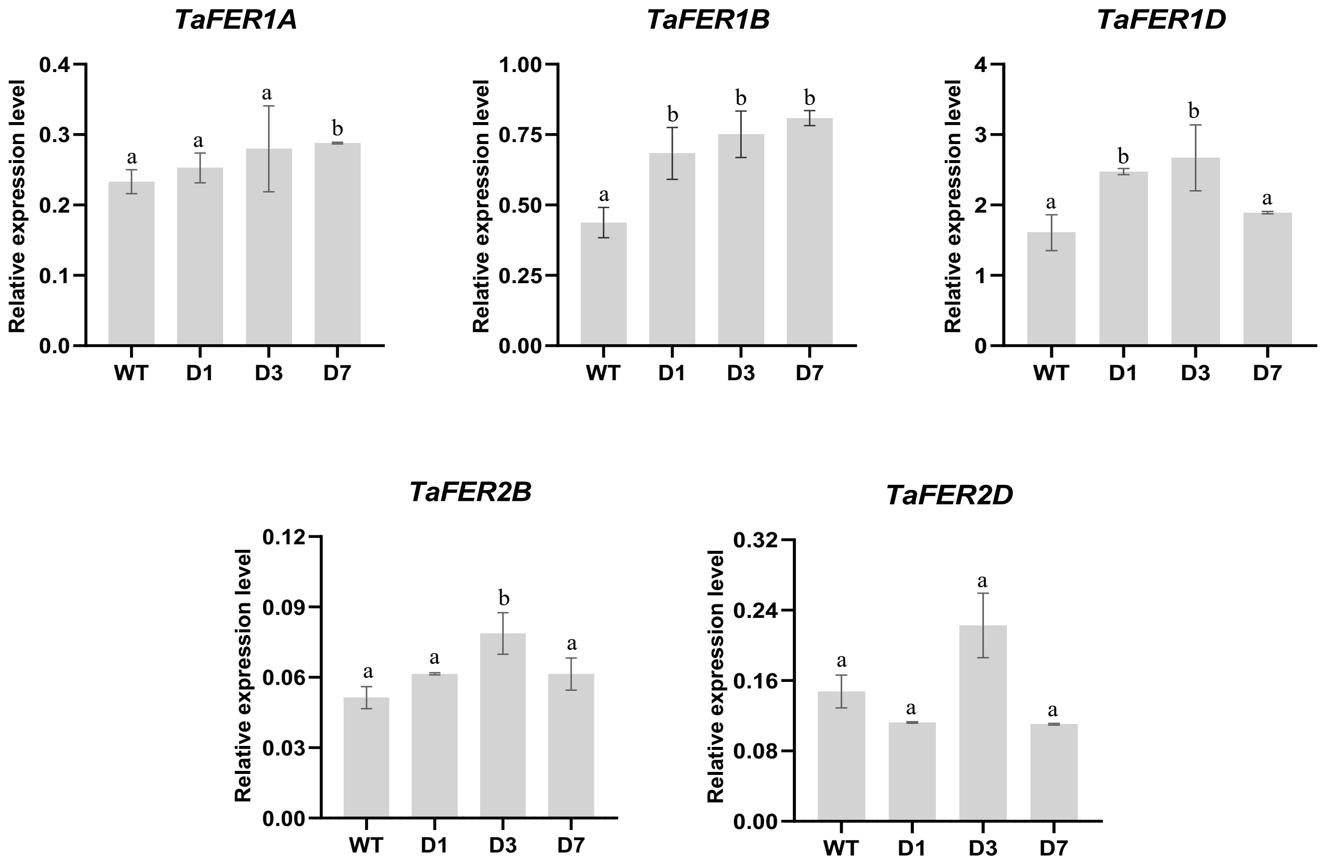


**Supplementary Figure S15**: qRT-PCR–based expression analysis of *Ferritin* genes in WT and TaHRZ1-edited (D1, D3, D7) seeds. Relative transcript levels of *TaFER1A*, *TaFER1B*, *TaFER1D* , *TaFER2B* and *TaFER2D* were analyzed in edited lines and WT seeds. Each bar represents the mean ± SD of three biological replicates. Gene expression levels were normalized to *TaARF1* as an internal reference. Different letters above the bars indicate statistically significant differences ($P < 0.05$), as determined by students t-test.
