## Supplementary Tables for "CRISPR/Cas9 editing of the wheat iron sensor *TaHRZ1* confirms its conserved role in iron homeostasis and allocation in grains"

**Supplementary Table S1:** List of the primers used in the current study

| **Primer for cloning** | | | |
| --- | --- | --- | --- |
| *TaHRZ2* | FP | pJET1.2 | ATGGCGCCGACGCCGATGGCCGGGGACGGCCCGA |
| *TaHRZ2* | RP | pJET1.2 | TTAATTCGACGTGGAACAGTCTGGTGCCTCCGTC |
| *TaHRZ1* | FP | pJET1.2 | ATGGCGACGCCCACGCCCATGGCCGGCGAGGGGA |
| *TaHRZ1* | RP | pJET1.2 | CTAGTTCGGGGTAGAACAATCTGCCGTATCAG |
| *TaHRZ2 gRNA1* | FP | JD633 | ACTTTACTGAAAAAGTTCCGTTGC |
| *TaHRZ2 gRNA1* | RP | JD633 | AAACGCAACGGAACTTTTTCAGTA |
| *TaHRZ2 gRNA2* | FP | JD633 | ACTTCACATTGAAGGCTGATACAT |
| *TaHRZ2 gRNA2* | RP | JD633 | AAACATGTATCAGCCTTCAATGTG |
| *TaHRZ1 gRNA1* | FP | JD633 | ACTTACACGTTCCAATAGCTTCAAT |
| *TaHRZ1 gRNA1* | RP | JD633 | AAACATTGAAGCTATTGGAACGTGT |
| *TaHRZ1 gRNA2* | FP | JD633 | ACTTAGGAATGTGCAAGTCTATCC |
| *TaHRZ1 gRNA2* | RP | JD633 | AAACGGATAGACTTGCACATTCCT |
| *TaBHLH404* | FP EcoR1 | pGADT7 | CCGGAATTCATGGCATCCCCGGAAGGATCAAACTGGG |
|  | RP Xho1 | | CCGCTCGAGCTATGCAACCGGAGGGCATGACTTGGGG |
| *TaBHLH405* | FP EcoR1 | pGADT7 | CCGGAATTCATGGCATCCCCGGAAGGATCAAACTGGGTA |
|  | RP Xho1 | | CCGCTCGAGTTATGCAACCGGAGGGCATGACTTGGGGTC |
| *TaBHLH406* | FP pGADT7  EcoR1 | | CCGGAATTCATGGCATCCCCGGAAGGATCAAACTGGGTA |
|  | Rp XhoI | | CCGCTCGAGTTATGCAACCGGAGGGCATGACTTGGGGTC |
| *TaBHLH407* | FP pGADT7  EcoRI | | CCGGAATTCATGTCTCTCCCTCCGACCGACGGCGGCGAC |
|  | RP Sma1 | | TCCCCCGGGTCACGCAACAGGCGGGCATGCTTCGCTGTC |
| *TaBHLH408* | FP EcoR1 | pGADT7 | CCGGAATTCATGTCTCTCCCTCCGACCGACGGCGGCGAC |
|  | RP Sma1 | | TCCCCCGGGCACGCAACAGGCGGGCATGCTTCGCTGTCC |
| *TaBHLH409* | FP EcoR1 | pGADT7 | CCGGAATTCATGTCTCTCCCTCCGACCGACGGCGGCGAC |
|  | RP Sma1 | | TCCCCCGGGTCACGCAACAGGCGGGCATGCTTCGCTGTC |
| *TaFIT* | FP EcoR1 | pGADT7 | CCGGAATTCATGGAGCACCACCAGCGGCTGCTGCATTTG |
|  | RP BamH1 | | CGCGGATCCTTAGCAGATCTCCGACGTGGCTTCTGGCCG |
| *TaHRZ2* | FP (EcoRI) | pGKBT7 | CCGGAATTCATGGCGCCGACGCCGATGGCCGGG |
| *TaHRZ2* | RP (SalI) |  | ACGCGTCGACTTAATTCGACGTGGAACAGTCTGGTGC |
| *TaHRZ1* | FP (Sma1) | pGKBT7 | TCCCCCGGGATGGCGACGCCCACGCCCATGGCAG |
| *TaHRZ1* | RP (Sal1) |  | ACGCGTCGACCTAGTTTGGGGTAGAACAATCTGCCGTATCA |
| tr.*TaHRZ1* | RP (SalI) | pGKBT7 | ACCGAATATTTGCTTCTCCTGGTCAC |
| tr.*TaHRZ2* | RP (Sal1) |  | TGATCTTGGTTGTGGTGATTTCTGCT |
| *TaHRZ1* | FP (Spe*I*) | pCAMBIA1302 | GGACTAGTATGGCGACGCCCACGCCCATGGCAGGCG |
| *TaHRZ1* | RP (SpeI) |  | GGACTAGTGTTTGGGGTAGAACAATCTGCCGTATCAGT |
| *TaHRZ2* | FP (BglII) | pCAMBIA1302 | GGAAGATCTATGGCGCCGACGCCGATGGCCGGGGACG |
| *TaHRZ2* | RP (Spe1) |  | GGACTAGTATTCGACGTGGAACAGTCTGGTGCCTCCGT |
| *TaHRZ1* gRNA |  | pRGEB32 | GGGGACAAGTTTGTACAAAAAAGCAGGCTATGAGGAATGTGCAAGTCTATCC |
| *TaHRZ*1 gRNA |  | pRGEB32 | GGGGACCACTTTGTACAAGAAAGCTGGGTTCAGGATAGACTTGCACATTCCT |
| *TaHRZ1* | Fp (SpeI) | BiFC | GGACTAGTATGGCGACGCCCACGCCCATGGCAGGCG |
|  | Rp (SalI) |  | ACGCGTCGACGTTTGGGGTAGAACAATCTGCCGTATCA |
| *TaBHLH406* | Fp (XbaI) | BiFC | TGCTCTAGAATGGCATCCCCGGAAGGATCAAACTGGG |
|  | Rp (XhoI) |  | CCGCTCGAGTGCAACCGGAGGGCATGACTTGGGGTCA |
| *TaBHLH408* | Fp (SpeI) | BiFC | GGACTAGTATGTCTCTCCCTCCGACCGACGGCGGCG |
|  | Rp (SmaI) |  | TCCCCCGGGCGCAACAGGCGGGCATGCTTCGCTGTCC |
| *TaFIT* | Fp (XbaI) | BiFC | TGCTCTAGAATGGAGCACCACCAGCGGCTGCTGCATT |
|  | Rp (BamHI) |  | CGCGGATCCGCAGATCTCCGACGTGGCTTCTGGCCGG |
| **SALK Genotyping Primers** | | | |
| bts-1 (SALK_016526) | FP | TGAATATGCTTGCTGTGTTCTTGC | |
|  | RP | CGTCTCAAAATCTGGTAACGG | |
| LB 1.3 |  | ATTTTGCCGATTTCGGAAC | |
| **Primers for qPCR** | | | |
| *TaHRZ2* | qrt-F | CCTGGCAGCACCTTCCTCTGAAA | |
| *TaHRZ2* | qrt-R | TGCTGGCCAATGAACATGGGCAG | |
| *TaHRZ1* | qrt-F | TCGCTCAGA GGATAAATCC | |
| *TaHRZ1* | qrt-R | TGCATCATTTCCTCGCTTGCCTGG | |
| *AtACTIN8* | FP | TCAGCACTTTCCAGCAGATG | |
| *AtACTIN8* | RP | CTGTGGACAATGCCTGGAC | |
| *AtFIT* | FP | TCGGTCTAGGACTTTGATCTCTG | |
| *AtFIT* | RP | TCTTGAACATACAACACTGCATCT | |
| *AtIRT* | FP | CTCTTTGCTTCCATCAAATGTTC | |
| *AtIRT* | RP | CCTAACGCTATTCCGAATGG | |
| *AtFRO2* | FP | TTCACCGTTCATGGTCTTTGTT | |
| *AtFRO2* | RP | GAGCTATCTCTCCGGCCAAATT | |
| *AtMYB* | FP | GGATAAACTATCTGAGACCGGACG | |
| *AtMYB* | RP | GAGATGCGTGTTCCACACGTTT | |
| *AtNAS4* | FP | TGTTCTTGGCTGCTCTTGTAGG | |
| *AtNAS4* | RP | CAAGGCTCAACGATTGGATAGA | |
| *AtPYE* | FP | CAGGACTTCCCATTTTCCAAG | |
| *AtPYE* | RP | CTTGTGTCTGGGGATCAGGTT | |
| *TaARF1* | FP | TGATAGGGAACGTGTTGTTGAGGC | |
| *TaARF1* | RP | AGCCAGTCAAGACCCTCGTACAAC | |
| *TaFIT* | FP | TGCGAGGGCGCGTCCCCGGACGAC | |
| *TaFIT* | RP | GCTCGTATAGCTTCTCCTTCATCCG | |
| *TabHLH404* | FP | TCCAAAGCATGCAGGGAGAAAGTGAGAAG | |
| *TabHLH404* | RP | CTTTAGCTCTCTAATCTTCTCTTGGA | |
| *TabHLH407* | FP | AGCTGAATGACAGGTTCCTTGAATTGGGTA | |
| *TabHLH407* | RP | TTTCTCTAGTTTCAGCTTCTGCTTCTC | |
| *TaIRO3* | FP | ATGGTCGCCATGCTGCCCG | |
| *TaIRO3* | RP | ATCATCCTCGGGGCCTTCTTCTTGC | |
| *TaNAS3* | FP | CGTCAATGTCACGTCGCT | |
| *TaNAS3* | RP | TGATAGCACACCTCGGTACA | |
| *TaFRO2* | FP | TACACCCACCAGCTCTACGTGGTC | |
| *TaFRO2* | RP | GAGAAGACTAGCTCCACCGTTCCG | |
| *TaIDEF1* | FP | ATGGCCGACGCGAACGGATCC | |
| *TaIDEF1* | RP | TCATTGTGATGGTGCTGAGGTGAAG | |
| *TaFer1-A* | Fp | GGTTTGTAGGGTGCTATGCTGCT | |
| *TaFer1-A* | Rp | CAAATCCCAAACACAGAGCAGTTC | |
| *TaFer1-B* | FP | GCTTGAAGGACACGGACACG | |
| *TaFer1-B* | RP | TATCACACCTACATCCTCCATTGC | |
| *TaFer1-D* | FP | TTTGCAGTGGGATTTGTAGGGTA | |
| *TaFer1-D* | RP | AGCGTGGC ACATCTCCCA | |
| *TaFer2-B* | FP | GTGGATCAGCCAACCACTGGT | |
| *TaFer2-B* | RP | GCAAACACCTTACACACCACTCTG | |
| *TaFer2-D* | Fp | GTGGATCAGCCAACCTCTGTG | |
| *TaFer2-D* | Rp | CGCAAACACCTTACACACCACTC | |
| **Primer for synthesizes sgRNA target for *in-vitro* validation** | | | |
| *TaHRZ2 gRNA1* | FP | CAGAATATGTTCCTGGCAGGTACC | |
| *TaHRZ2 gRNA1* | RP | GCTATGTGCTCTGTAAAGACCCC | |
| *TaHRZ2 gRNA2* | FP | GTAGGTTGTATTTGCATGGATAGGA | |
| *TaHRZ2 gRNA2* | RP | AGGCATAGAACAAACTATTAATTAAA | |
| *TaHRZ1 gRNA1* | FP | GGCAGGAGCAAAATCAACTGCAGC | |
| *TaHRZ1 gRNA1* | RP | CCAGATGAAGAGCAACATCTGCAC | |
| *TaHRZ1 gRNA2* | FP | CATTCTGCCCATGTGCTTCACGTATC | |
| *TaHRZ1 gRNA2* | RP | CTTTTGCTGAGCAGCTATCCAGCGG | |
| **Screening Primers** | | | |
| grna | FP | TGAACCAGCCACATACTGAA | |
|  | RP | TCAGCGCTTAATCAAGAAGAACAGA | |
| CAS9 | FP | GGCCTGTTCGGGAACCTCATCGCT | |
| CAS9 | RP | AGGGTCAGATCCTGATGGTGCTCG | |
| JD633 | FP | CCGGTGACGGACGACCAAG | |
| JD633 | RP | AAACACTGATAGTTTAAACTGAAGGCG | |
| HYG | FP | TACGCCCGACAGTCCCGGCTCCGG | |
| HYG | RP | TCGGACCGCAAGGAATCGGTCAAT | |

**Supplementary Table S2:** Transformation efficiency of different cultivars after transformation with pRGEB32 and JD633

| **Constructs** | **Batch number** | **No. of immature embryos** | **Callus** | **Callius surviving screening** | **Callus with shoots** | **Regeneration efficiency (%)** |
| --- | --- | --- | --- | --- | --- | --- |
| **cv. Fielder** | | | | | | |
| *pRGEB32* | 1  2  3 | 332  338  251 | 104  117  85 | 31  34 | 3  7  4 | 1.4 |
| *GRF4/GIF1* | 1  2  3 | 318  328  315 | 220  257  292 | 69  78  32 | 26  37  22 | 8.8 |
| **cv. C306** | | | | | | |
| *pRGEB32* | 1  2 | 123  148 | 34  28 | 27  18 | 1  1 | 0.73 |
| *GRF4/GIF1* | 1  2  3 | 187  156  268 611 | 68  49  80 | 36  31  21 | 12  14  13 | 6.38 |
